# Beyond Proxies: Towards ecophysiological indicators of drought resistance for forest management

**DOI:** 10.1101/2025.02.06.636586

**Authors:** Alice Copie, Caroline Scotti-Saintagne, François Lefèvre, Hervé Cochard, Sylvain Delzon, Pierre-Jean Dumas, Damien Gounelle, Stephane Herbette, Florence Jean, Nicolas Mariotte, Maurizio Mencuccini, Ivan Scotti, Nicolas Martin-StPaul

**Affiliations:** INRAE-URFM, Domaine Saint Paul - 228 route de l’Áerodrome - Site Agroparc - CS 40509, Avignon Cedex 9, 84914, France; INRAE, PIAF, Université Clermont Auvergne, Aubière, 63178, France; INRAE, UMR BIOGECO, Université de Bordeaux, Pessac, 33318, France; CREAF, Bellaterra, Barcelona, 08193, Spain

**Keywords:** *Keywords*: Hydraulic Failure, Integrated traits, Model, *Abies*, Trees

## Abstract

As drought-induced mortality increases globally in forest biomes, it becomes necessary for foresters to have access to reliable predictors of species vulnerability to drought and mortality risk under different climatic scenarios. On the one hand, there exist several “operational” indicators of drought resistance, which are based on observations, expert knowledge and species bioclimate (*e.g.* Ellenberg, Rameau, ClimEssences). On the other hand, as traits can now be measured at high throughput, functional traits (such as plant hydraulic traits) have been increasingly used to assess species’ vulnerability to hydraulic failure, a key process of tree mortality under drought. However, this mechanistic approach has never been compared to the operational approach. In this study, we review if indicators commonly used by foresters provide information on *Abies* species’ vulnerability to hydraulic failure. We measured a set of traits in a common garden experiment of closely related Mediterranean *Abies* species. These traits were used as inputs to the SurEau model to compute a single indicator of vulnerability (Time to Hydraulic Failure - *THF*) and to assess mortality risk in future climate. We found that among circum-Mediterranean firs, a high *THF* was mainly due to high water losses after stomatal closure. Some operational indicators are good proxies of *THF*, however they are not available for all species, reduce a species to a single value and have the same limitations as species distribution models. We argue that the ecophysiological approach could help foresters in species selection and in estimating the risk faced by forest tree species in a changing climate. While accounting for the variability of traits, hydraulic models can be forced with different climatic scenarios allowing hydraulic failure risk assessment by the end of the century.

**Highlights:**

- Integrating ecophysiological traits in a process-based model allows the computation of an accurate indicator of vulnerability: the Time to Hydraulic Failure (*THF*).
- For *Abies* species, ”operational” indicators, *i.e.* indicators used by foresters and land managers, are mutually consistent and can be simplified proxies of the vulnerability to hydraulic failure.
- More research on intra-specific variability is needed to assess the evolutionary potential of Euro-Mediterranean *Abies* species.

## 1. Introduction

Recent reports that the European forest carbon sink is shrinking have again emphasised the need for adaptive management and protection of forest ecosystems (Korosuo et al., 2023; Chakraborty et al., 2024). Although the strong selection pressures exerted by climate on populations contribute to new local adaptations (Aitken and Bemmels, 2016), the fear that the speed of climate change may exceed the natural adaptive capacities of tree populations (Shaw and Etterson, 2012), raises major concerns about the persistence of numerous species (Kremer et al., 2012; Quigley et al., 2019; Radchuk et al., 2019; Martin et al., 2023). For this reason, new ecosystem management strategies that involve human intervention are being proposed (Peterson St-Laurent et al., 2018; Gaitán-Espitia and Hobday, 2021), including the introduction of non-local provenances or species (Aitken and Whitlock, 2013). As major forest die-backs have been linked to the increasing frequency and duration of drought events (Allen et al., 2015), possibilities of replenishing existing species populations with neighbouring species having traits conferring greater resistance to drought and that could eventually hybridise are actively discussed, particularly in the context of the European Green Deal, which aims to plant 3 billion trees by 2030. However, accurate knowledge of species ecology is required, both in local species and in closely related taxa (Kurz et al., 2023).

Currently, there are different ways to assess drought resistance of tree species. Often, this relies on the span of bioclimatic variables (temperature, precipitation) observed in a species’ range, with the inherent limitation that the data reflect the realised distribution and not the potential distribution of the species (Jiménez-Valverde et al., 2008; Ay et al., 2017; Jourńe et al., 2020). Drought resistance can also be assessed through greenhouse drought experiments (Valladares and Śanchez-Gómez, 2006; Urli et al., 2013) or by studying tree growth in relation to climate (Lloret et al., 2011; Fuchs et al., 2021). For the purpose of species comparisons, Niinemets and Valladares (2006) have summarised a species’ drought resistance to a single number, using large-scale reviews of data on 806 species. Land managers, and in particular foresters, use their own single-digit indicators of drought resistance. In France, the widely used ”adult tree resistance to severe drought” of ClimEssences (RMT AFORCE, 2021) integrates mortality data gathered on local forests and existing literature on forest tree species. Rameau’s Flora (Rameau et al., 2016) or Ellenberg’s (Ellenberg and Leuschner, 2010) indicators, also in use among foresters, are based on empirical knowledge of species functional ecology. However, these indicators summarise a species to one value, are not available for all species, do not take into account plasticity nor the genetic variability within species. Furthermore, these indicators are based on empirical knowledge based on past and current climatic conditions.

Using functional traits to assess the drought response of forest tree species is a complementary approach to previously mentioned indicators. A wide variety of traits are used for this purpose. On the one hand, traits related to growth potential and water-use strategies, such as Huber Value or wood density, have been used as proxies of tree survival under drought (Valladares and Śanchez-Gómez, 2006; Chave et al., 2009; Matías et al., 2012; Lopez-Iglesias et al., 2014; O’Brien et al., 2017). On the other hand, ecophysiological plant hydraulic traits have recently been receiving high attention for assessing species resistance to drought (e.g. Kunert and Tomaskova (2020); Skelton et al. (2021)). Indeed, hydraulic failure – a physiological process related to increased tension (water potential) in the plant’s conductive tissue (xylem) and potentially leading to loss of water transport – has been identified as one of the main mechanisms involved in tree mortality under drought (Adams et al., 2017; Sanchez-Martinez et al., 2023).

To limit hydraulic failure, trees have developed two different strategies: tolerance and avoidance. Tolerant trees are able to sustain high tension in their tissue to maintain gas exchange and growth longer during drought, while others avoid experiencing high tension by limiting water losses through growth reduction and stomatal closure (Kramer, 1983; Jones, 1992; Delzon, 2015). Encompassing these two strategies, multiple traits articulate together to shape the diversity of drought resistance mechanisms among tree species and define the sequence of plant dehydration during drought (Martin-StPaul et al., 2017; Choat et al., 2018). One of the first events occurring when a plant dehydrates is the progressive loss of cell turgor (Choat et al., 2018), which is tightly correlated with stomatal closure and the loss of productivity (Brodribb and Holbrook, 2003; Martin-StPaul et al., 2017). Turgor loss point (Π*_T LP_*) has thus widely been used as a proxy of drought resistance (Kunert and Tomaskova, 2020; Sjöman et al., 2018). In spite of stomatal closure, plants continue to lose water through poorly closed stomata or through cuticles (Choat et al., 2018). This rate of water losses after stomatal closure is referred to as minimal or residual stomatal conductance *g_min_* (Duursma et al., 2019). This trait is increasingly recognised as important in driving water use strategies (Lanning et al., 2020), and it deeply contributes to hydraulic failure risk (Duursma et al., 2019). As water potential continues to decrease with residual losses, xylem cavitation can occur (Choat et al., 2018). Excessive embolism is linked to the loss of plant hydraulic conductivity and thereby to hydraulic failure. The water potential causing 50% loss of hydraulic conductance (*P*_50_), is a key trait involved in drought tolerance, that has been the focus of large research efforts over the last decades (Maherali et al., 2004; Choat et al., 2012; Lens et al., 2016).

More recently, in order to take into account both xylem resistance to embolism and water loss regulation, researchers have used integrated indicators such as the Stomatal Safety Margin (*SSM*) or the Stomatal Margin Retention Index (*SMRI*) (Blackman et al., 2016; Martin-StPaul et al., 2017; Petek-Petrik et al., 2023). In a similar purpose, functional models are being increasingly used to estimate vulnerability to hydraulic failure (Cochard et al., 2021; Ruffault et al., 2022).

Here, we define the ”operational” approach as one that tends to summarise empirical observations on species ecology with a single figure, as opposed to the ”mechanistic” approach using functional and ecophysiological traits. The operational indicators we consider are seven mean species bioclimatic variables (Fick and Hijmans, 2017; Zomer et al., 2022) and four drought resistance indicators (Aussenac, 1980; Rameau et al., 2016; RMT AFORCE, 2021; Tichý et al., 2023). For the mechanistic approach, we consider eight functional traits and the output of the hydraulic model SurEau (Time to Hydraulic Failure). The operational and the mechanistic approach have never, to our knowledge, been compared. The *Abies* genus of the circumMediterranean area provides an interesting framework for the comparison of indicators. Indeed, current threats and observed intensive dieback (Ĺopez-Tirado et al., 2023; Piedallu et al., 2023) to the widely represented European species *Abies alba* make other fir species of the *Abies* and *Piceaster* sections potential candidates for adaptation measures. Study of firs’ past evolution suggest that they diverged during the late Oligocene-Early miocene, leading to two *Abies* sections: *Piceaster* and *Abies*, with permeable boundaries (Balao et al., 2020). Euro-Mediterranean firs hybridise easily (Krajmerová et al., 2016) and occur over a wide geographic area including southern, drought-prone environments (Linares, 2011).

Studying closely related *Abies* species, we investigated how operational indicators inform us on the vulnerability to hydraulic failure, and how the ecophysiological approach, used to quantify drought resistance, could benefit forest management. First, we computed a synthetic representation of operational indicators using principal component analysis. Second, we compared the different functional traits measured in eight *Abies* species. We then used these traits as inputs to the SurEau process-based model (Cochard et al., 2021; Ruffault et al., 2022) to compute a single indicator of vulnerability to hydraulic failure: Time to Hydraulic Failure (*THF*), and assessed how each trait influenced *THF* . Third, we compared the PCA-based synthetic representation of operational indicators to *THF* and, last, we explored an applied use of the SurEau model for risk assessment by the end of the century under different climatic projections.

## 2. Material and methods

### 2.1. Experimental setup used for physiological measurements

*Abies* species are found in mountainous regions at altitudes ranging from 300 to 2200 meters. *A. bornmuelleriana* is the only fir found at see level (euforgen.org, last accessed in August 2023). Sampled Mediterranean firs were representative of the *Abies* genus in the circum-Mediterranean area. Other firs exist in the Mediterranean area such as *A. nebrodensis, A. equi-trojani* and *A. tazaonata*. However, *A. nebrodensis* is highly endemic (only 24 living individuals in its natural range area), while *A. equi-trojani* and *A. tazaonata* are respectively sub-species of *A. nordmanniana* and *A. pinsapo*. We sampled trees from an experimental plot located in ”Le Treps” in the Maures forest (Mediterranean forest, South of France, Longitude: 6.373653, Latitude: 43.262764). Soil is sandy and loamy and bedrock is composed of silica and shale. The plantation grows on flat ground, altitude is 600 meters, mean annual temperature was 15°C, and mean annual rainfall was 800 millimetres, for the 1970-2019 period; maximum and minimum recorded temperatures were respectively 40°C (July 1982) and -9°C (January 2002). In 1970 and 1977 seedlings (five and four years old) were transplanted from the nurseries on two different sites within the same forest, one kilometre apart. Trees were spaced two meters apart. Both setups are described in Fady et al. (2023, 2024) and summarised in Appendix A. We sampled eighty trees from eight *Abies* species for hydraulic traits measurements. We selected ten trees per species from one or two provenances. Seven species originated from the Mediterranean basin, from East Türkiye to Morocco. *A. pinsapo marocana* and *A. numidica* belong to the *Piceaster* section. *A. borisii-regis, A. bornmuelleriana, A. cephalonica, A. cilicica* and *A. nordmanniana* belong to the *Abies* section. The eighth species, *A. concolor*, originated from Colorado (USA) and belongs to the *Grandis* section. The geographic origins of the provenances selected for each species are presented in figure 1 and sampling is summarised in supplementary table A.1.

**Figure 1:**
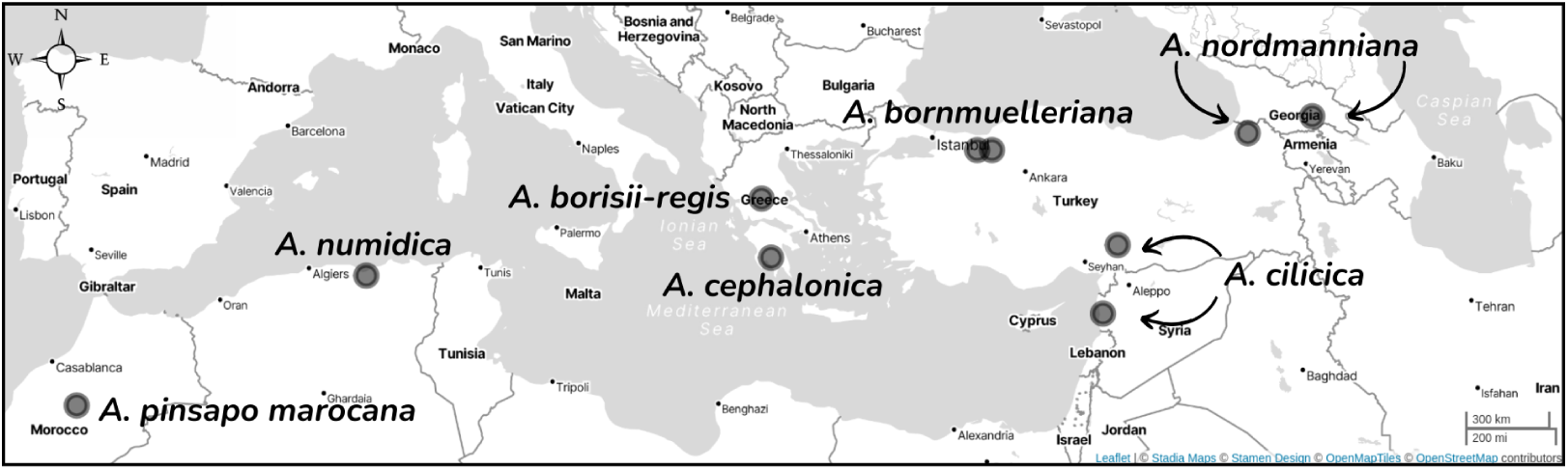
Origin of the seven Mediterranean *Abies* fir species planted on the experimental site of ”Le Treps”. For *A. bornmuelleriana*, *A. cilicica* and *A. nordmanniana*, two provenances were sampled. The American fir, *A. concolor*, originating from Colorado was also sampled.

*A. alba* did not survive to the climatic conditions in Le Treps. Therefore, to compare this key species to our sampled Mediterranean species, we collected values of ecophysiological traits from the literature and values measured at another experimental site at the Mont Ventoux mountain (France). The values used in the analyses and simulations are presented in supplementary table A.2.

### 2.2. Bioclimatic variables

For each species, we extracted the Aridity Index (ratio between Mean Annual Precipitation and Mean Annual Reference Evapotranspiration) and potential evapotranspiration (ET0) (Zomer et al., 2022) . We also extracted five drought-related bioclimatic variables from the WordClim database (Fick and Hijmans, 2017): annual mean temperature (bio1), mean temperature of warmest quarter (bio10), annual precipitation (bio12), precipitation seasonality or coefficient of variation (bio15), precipitation of warmest quarter (bio18). We extracted values based on species distribution range areas defined by the European Forest Genetic Resources Programme (euforgen.org, last accessed in August 2023), based on expert knowledge. For *A. concolor*, we used Global Biodiversity Information Facility points (gbif.org, last accessed in August 2023), keeping only the points located in the United States. For all species, we used 30 arcsecond resolution, except for *A. alba* for which we used 2.5 arcminute resolution because this species has the largest range area. We estimated that a sufficient number of points was obtained with 2.5 arcminute resolution. Using R, we computed 0.05 quantiles, average values and 0.95 quantiles from the points at the intersection between the variable’s grid and the distribution area of the species. For *A. concolor*, we extracted values of variables from occurrence points in the United States. Additionally, we extracted bioclimatic variables of original provenance locations and of Le Treps experimental site.

### 2.3. Drought resistance indicators

We collected information on four drought resistance indicators. Drought resistance of Mediterranean firs was previously assessed by the pioneering work of Aussenac in 1980 who measured resistance to desiccation of excised branches. From his measurements, he deduced a drought resistance index for seven circum-Mediterranean firs (Aussenac, 1980). In France, decision makers and foresters now often use the ”adult tree’s resistance to severe droughts” from ClimEssences website (RMT AFORCE, 2021). This indicator is based on available literature and recent observations of drought-related mortality events), such as mortality data from 2003 (*A. alba*, *A. bornmuelleriana* and *A. concolor*), 2006 (*A. concolor* and *A. bornmuelleriana*) and 2017 (*A. cephalonica*). For exotic species for which no mortality data were available, only references from literature were used (*A. nordmanniana*, *A. cilicica*, *A. numidica* and *A. pinsapo*). No data was available for *Abies borisii-regis* on ClimEssences. We also collected two other indicators: Rameau’s Flora Xericity indicator (Rameau et al., 2016) and Ellenberg’s resistance to drought (Ellenberg and Leuschner, 2010). Both indicators are computed according to ”expert-based rankings of plant species according to their ecological optima on main environmental gradients” (Tichý et al., 2023). The four drought resistance indicators were not available for all species. Eight species were available for Rameau and ClimEssences, seven for Aussenac and five for Ellenberg. We grouped mean species bioclimatic variables and drought resistance indicators under the term of ”operational” indicators.

### 2.4. Basic hydraulic traits

We measured four sets of traits related to the drought performance of trees, including: (i) resistance of the xylem to cavitation from the vulnerability curve (potential inducing 50% loss of conductivity, slope at the inflection point); (ii) leaf traits derived from the pressure volume curves (turgor loss point, osmotic potential at full turgor, modulus of elasticity); (iii) minimal water losses after stomatal closure (minimal conductance); and (iv) we computed two integrated hydraulic traits.

#### 2.4.1. Vulnerability Curves

In November 2019, we sampled one approximately 50 cm-long light-exposed branch per tree on ten trees per species (eighty branches). Immediately after collecting, we cut the tips of the branches and placed them under water; then we trimmed their proximal end while still immersed, to eliminate potentially embolized conduits and we removed the foliage. We stored the branches in plastic bags saturated with humidity to transfer them to the high-throughput phenotyping platform (Phenobois, INRAE University of Bordeaux). Prior to measurement, we again immersed branches in water, removed the bark and trimmed them to 28 cm to suit the size of the rotor centrifuge. We assessed vulnerability to cavitation using the Cavitron technique which we summarise hereafter (Cochard, 2002). Using a rotor centrifuge, a branch is spun to decrease its water potential while estimating water flow conductance. Measurements are performed at increasing angular speed, generating increasingly negative water potentials, which allows drawing a vulnerability curve. Using the Cavisoft algorithm (Cavisoft v.4.0, University of Bor-deaux), we calculated the percentage of lost conductivity (*PLC*) of samples as: 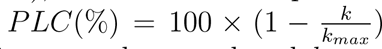, where *k* is the conductivity recorded at a given angular speed and *k_max_* the maximal conductivity recorded at the lowest angular speed. We constructed xylem vulnerability curves by plotting *PLC* against pressure and fitting the following sigmoid function: 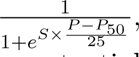 with *S* and *P*_50_ the two key traits to be estimated. *P*_50_ is the water potential inducing 50% loss of hydraulic conductivity and *S* is the slope of the curve at the inflection point.

#### 2.4.2. Pressure Volume curves

For five of the ten branches per species used for the vulnerability to cavitation, we collected apical twigs for pressure-volume curve measurements. Once collected, we bagged and placed the twigs in a cooler (4°C) until reaching the laboratory. Once in the lab, we trimmed the proximal end of each twig under water with a razor blade, then we put them overnight in a cold chamber in distilled water to re-hydrate. We used pressure–volume curves (hereafter PV curves) to characterise different foliar traits related to the maintenance of leaf turgor and hydration (Tyree and Hammel, 1972; Bartlett et al., 2012). We established PV curves using the bench drying method proposed by Hinckley et al. (1980) and recently applied by Limousin et al. (2022). We measured the weight and the water potential of the twigs during dehydration using a precision balance (FS-220, resolution 0.1 mg) and a Scholander pressure chamber (PMS instrument), respectively. To correct for oversaturation of rehydrated twigs, we removed the first measurement of water potential and weight, and we extrapolated full turgor weight from the regression between twig weight and water potential before the turgor loss point. We obtained the osmotic potential at full turgor (*π*_0_) and at the turgor loss point (Π*_T LP_*), the relative water content at turgor loss point (*RWC_T LP_*), the apoplasmic water fraction at full turgor (*a_f_*), and the bulk modulus of tissue elasticity (*ɛ*) by plotting the inverse of water potential (− <u>^1^</u>, in MPa) against twig water saturation deficit (1 − *RWC*, in %). Because PV curves were measured on a subsample of individuals (compared to the other traits), we used average species level parameter values (Π*_T LP_*, *π*_0_, *ɛ*, *LDMC*, *a_f_* and *LMA*) for the computation of integrated traits and modelling exercises.

#### 2.4.3. Minimum conductance

For eight trees per species, among the ten previous ones, we computed minimum conductance (*g_min_*) from water losses measured under controlled condition using the DroughtBox chamber (Billon et al., 2020), which works as follows. Tree branches are attached to strain gauges in a climatically controlled chamber. Gauge tension is recorded over time and then converted to weight. Temperature and relative humidity are measured continuously and automatically adjusted to keep them constant. The droughtBox can host eight branches at once. We sampled full-light branches in spring 2023 (between the 20th of March and the 18th of April). Immediately after cutting, we trimmed their proximal ends under water, wrapped the samples in wet paper and stored them in a sealed plastic bag, to prevent embolism and leaf dehydration. We performed drought box measurements the following week. We rehydrated the samples for a minimum of 12 hours: we again trimmed their proximal ends under water and soaked the samples overnight. The following day, we trimmed branches above the wet end. We measured water potentials, weights, lengths and diameters at the beginning of the experiment. We sealed proximal twig ends with paraffin. We hanged the samples in the drought box chamber. We set the box to 30°C and 40% relative humidity. After 2.5 days of measurement, we weighted the samples and oven-dried them at 60°C for 72 hours. To identify the plateau phase, we plotted water content per unit of projected leaf area as a function of time, according to equation 1.

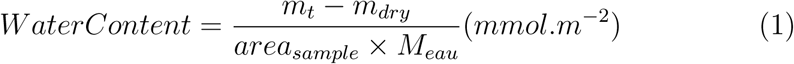

At the beginning of the measurement, water content drops rapidly as the samples lose great amounts of water due to open stomata. After an inflection point (i.e. stomata are closed), a stationary regime is reached. *g_min_* is taken as the slope of the curve during this stationary regime (equation 2).

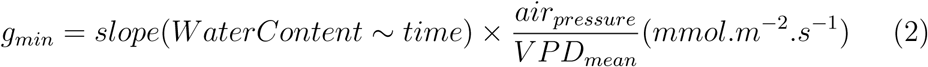

### 2.5. Integrated hydraulic traits

We computed two integrated traits: Stomatal Safety Margin (*SSM*) and Stomatal Margin Retention Index (*SMRI*). *SSM* is defined by MartinStPaul et al. (2017) as the difference between the water potential inducing stomatal closure and *ψ*_50_. Similarly to Martin-StPaul et al. (2017), we used the turgor loss point (*ψ_tlp_*) obtained from pressure volume curves described above, as a proxy of *ψ_close_*. Therefore, we had the following equation : *SSM* = Ψ*_tlp_* − Ψ_50_. We calculated individuals *SSM* using average Ψ*_tlp_* per species and individual values for Ψ_50_. *SMRI*, which also includes the minimum conductance (*g_min_*) is an indicator of the time required to cross the Stomatal Safety Margin and is defined by Petek-Petrik et al. (2023) as: *SMRI* = *SSM/g_min_*.

### 2.6. Other traits

We also measured the following traits: Specific Leaf Area (*SLA*), Huber Value (*HV*) and succulence.

We measured *SLA* using additional branch samples collected together with drought box samples and stored in a similar way until reaching the laboratory. Branches ranged from 14 to 80 g. We randomly collected between 50 and 60 leaves on the sample and scanned them with an Epson® scanner. We analysed images with Grass® and Qgis® to extract projected leaf areas. We measured dry weight of the scanned leaves after 72h in a 60°C oven. We calculated *SLA* as the ratio between projected leaf area and leaves dry weight.

*HV*, defined as the ratio of a plant’s water supply (sapwood area) over its transpiration area (leaf area), is an indicator of a plant’s water balance. We calculated *HV* on *SLA* samples. We measured sapwood area based on the diameter under bark of the fresh branch, assuming no heartwood. We calculated leaf area as the ratio between the previously measured *SLA* and the leaf dry weight of the sample.

To account for the tree’s water reserves, we computed succulence as the ratio of a twig’s water content over its projected leaf area. We used drought box samples for this calculation.

### 2.7. Modelling approach

#### 2.7.1. Model description

We explored how *Abies* physiological parameters interact to shape the vulnerability and the risk to hydraulic failure using the soil plant hydraulic model SurEau (Martin-StPaul et al., 2017; Cochard et al., 2021; Ruffault et al., 2022; Mas et al., 2024; Moreno et al., 2024). For this work, the SurEauEcos version 2.0 was used (Ruffault et al., 2022). SurEau simulates water fluxes from the soil to the atmosphere, the water potential in different plants organs, and thus the evolution of embolism in the different organs, stem and leaves. At each time step (typically 30 minutes), the model computes leaf stomatal and cuticular transpiration as the product between leaf-to-air VPD and stomatal and cuticular conductance. Then, stomatal and cuticular fluxes are used to compute plant water potential in the different plant compartments, while accounting for symplasmic capacitance and hydraulic conductance losses due to xylem embolism. Stomatal closure is regulated in a feedback manner, as a function of leaf water potential, through empirical relationships (Klein, 2014). Soil water potential and soil hydraulic conductance are computed from soil water content using Van Genuchten moisture retention curves (Van Genuchten, 1980). We parametrised the model with measurable plant traits. This includes (i) the traits that determine water potential for a given soil and atmospheric drought, such as plant hydraulic conductance, stomata closure sensitivity to water potential, and cuticular conductance; (ii) the traits that determine the responses of plant function to water potential such as plant vulnerability to cavitation, or pressure-volume curves that drive turgor loss and (iii), leaf traits that account for differences in water reserves and leaf morphology (leaf dry matter content and leaf mass area).

#### 2.7.2. Estimation of species vulnerability under standardised climatic conditions and at the individual level

We ran the model by setting constant climatic conditions from day to day, using weather data typical of a Mediterranean summer and we set the rain to 0. We stopped the simulation when the leaves reached 100% of embolism. We took the number of days until this date as the indicator of species vulnerability to hydraulic failure: Time to Hydraulic Failure (*THF*). For the eight *Abies* species, we used the parameters presented in figure 2 as variables. We carried out simulations assuming no segmentation of the vulnerability to cavitation and therefore using the same measured value for the leaf and stem. We evaluated the empirical response of stomata to water potential from the turgor loss point obtained from the pressure-volume curves as in Martin-StPaul et al. (2017). We set stand parameters (such as leaf area index) and soil properties identically between species in order to test only for the effect of hydraulic traits. We computed *THF* for each individual by considering the inter-individual variability of traits for *g_min_*, *P*_50_ and *S* of the vulnerability curve to cavitation. The model has been shown to be very sensitive to these traits (Cochard et al., 2021; Ruffault et al., 2022) and they were measured on a larger number of individuals. The traits measured on the pressure volume curve samples, including the following parameters: Ψ*_gs_*_12_, Ψ*_gs_*_88_, *ɛ*, *π*_0_, *a_f_*, LDMC, LMA, *gs_max_* were set to the mean species values. For *Abies alba*, we couldn’t estimate variability as this species died on our experimental site. Consequently, we ran simulations using mean trait values extracted from the literature (see supplementary table A.2).

**Figure 2:**
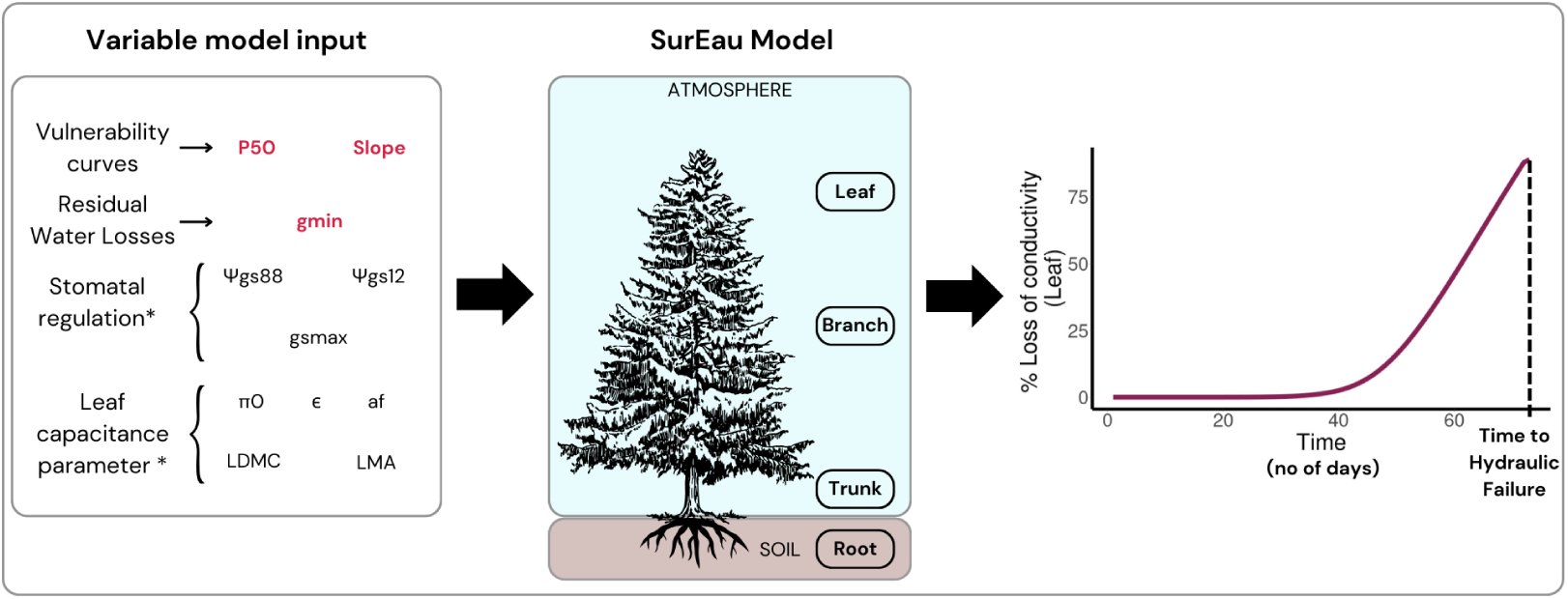
Conceptual framework of the modelling approach using the SurEau model. In red, individual values for vulnerability curves parameters and residual water losses were used as inputs: water potential causing 50% cavitation in the xylem (*P*_50_), slope of the vulnerability curve at *P*_50_ and minimum conductance (*g_min_*). Other traits were set to mean species values: Water potential causing 88% (respectively 12%) stomatal closure(Ψ*_gs_*_88_ and Ψ*_gs_*_12_), maximal stomatal conductance (*gs_max_*), osmotic potential at full turgor (*π*_0_), elasticity modulus (*ɛ*), apoplasmic fraction (*a_f_*), Leaf Dry Matter Content (*LDMC*) and Leaf Mass per Area (*LMA*). (*) Traits were derived from the leaf pressure volume curves.

#### 2.7.3. Risk assessment under climate change scenario for the different average fir species

Secondly, we aimed to estimate the risk of hydraulic failure at Le Treps site at the end of the 21st century using climate projection with three different warming scenarios. In order to reduce the computational load, we used mean trait values for each species (vulnerability). We forced SurEau with climate projections at Le Treps site (hazard) and used site measurements of LAI and extractable soil water (exposure), therefore combining the tree components of risk as defined by the Intergovernmental Panel on Climate Change (IPCC, 2018). We used down-scaled regionalised projections of *IPCC*’s Representative Concentration Pathways (RCP), provided by the DRIAS (*Drias^lesfutursduclimat^*) project (drias-climat.fr). The Global Climate Model (GCM) used by the DRIAS were produced within the framework of the 5th phase of the Coupled Model Intercomparison Project international exercise (CMIP5) which served as the basis for the 5th *IPCC* report (IPCC, 2014). The Regional Climate Model (RCM) were produced by the EuroCordex project (Jacob et al., 2014). The DRIAS proposes twelve pairs of GCM/RCM declined over two or three RCPs: we chose the EC/RACMO simulation. This simulation is representative of the median precipitations and temperatures of the twelve simulations and is declined over three scenarios. RCP2.6 corresponds to a virtuous world, sober in greenhouse gas emissions. At the opposite, RCP8.5 corresponds to a world with no climate regulation policies, leading to an increase of 5°C by the end of the century. RCP4.5 is intermediate between the two previous scenarios (Soubeyroux et al., 2020). We computed the risk as the probability of reaching more than 50 % yearly maximum percent loss of conductivity for each species in each RCP during the 2070-2100 time period.

### 2.8. Statistical analysis

To represent the variations among operational indicators and their coordination, we performed a Principal Component Analysis (PCA) with the four drought resistance indicators and the seven bioclimatic variables presented in sections 2.2 and 2.3. We set missing values of drought indicators to the mean value of the indicator so as not to influence its representation on the PCA axes. We extracted species coordinates on the two first dimensions of the PCA (PCA-dim1 and PCA-dim2) and used them for regression analysis with *THF* .

We performed a statistical analysis to estimate inter-specific differences of measured traits, integrated traits and *THF* . We used Levene’s test to check for homogeneity of variances between species. We visually checked data for non-normality using qqplots representations (see supplementary figure C.1). As Levene’s test was not significant, we performed an analysis of variance (ANOVA) using stats::aov function in R and we used Tukey’s post hoc test to analyse pairwise differences between species.

We related *THF* values to hydraulic traits using regression analysis to determine which trait best explains *THF* . We carried out the first analysis using average species values, thus including *Abies alba*. We then carried out the second analysis using the individual values of the different measured species. Similarly, to determine whether operational indicators are proxies of *THF*, we performed a regression analysis using PCA-dim1, PCA-dim2 and operational indicators as explanatory variables of mean *THF* .

To compare our results to other studies linking physiological traits and ecological or climatic preferences of species (Choat et al., 2012; Lens et al., 2016; Larter et al., 2017), we tested Pearson correlations between traits and operational indicators. Results are given in supplementary material C.3.

## 3. Results

### 3.1. Operational indicators

Species with large range areas such as *A. nordmanniana* showed high variability of bioclimatic variables (*e.g.* mean annual precipitation ranged from 531 to 1583 mm) while species with small range areas such as *A. numidica* showed low variability of bioclimatic variables (*e.g.* mean annual precipitation ranged from 827 to 1023 mm). The climatic conditions of Le Treps are within the range of climatic conditions for most of the species studied. For *A. alba*, 4/7 variables in le Treps were out of the species’ range (0/7 for *A. borisii-regis*, 3/7 for *A. bornmuelleriana*, 1/7 for *A. cephalonica*, 5/7 for *A. cilicica*, 3/7 for *A. concolor*, 1/7 for *A. nordmanniana*, 3/7 for *A. numidica* and 4/7 for *A. pinsapo*). Some provenances were beyond the species’ distribution ranges defined by EUFORGEN. *A. cilicica*’s sampled seeds originated from a hotter area than the species’ range values and precipitations were higher for one of the two sampled provenances. For *A. concolor*, sampled seeds originated from a drier and hotter area than mean species values, with almost no precipitations during the warmest quarter. *A. nordmanniana* originated from the hot and dry end of the distribution area. *A. pinsapo*’s seeds originated from a drier area than species’ range values. Bioclimatic variables for the entire species’ range area and the sampled seed location are presented in supplementary table B.5 along with the experimental site values.

Values of drought resistance indicators are presented in supplementary table B.6. All indicators agree on *A. alba* being less resistant to drought than other firs. However, drought resistance indicators disagreed on the classification of other species. Aussenac classified *A. concolor* as highly resistant to drought (A) whereas this species was classified as poorly resistant (C) by ClimEssences and Rameau. Similarly, Aussenac and Ellenberg classified *A. nordmanniana* as poorly resistant to drought (C, 5) whereas ClimEssences and Rameau classified it as resistant to drought (B, xerophile).

We analysed relations between the four drought resistance indicators (Rameau, Aussenac, Ellenberg and ClimEssences) and bioclimatic variables for nine *Abies* species using principal component analysis (figure 3). The first axis of the PCA (dim1) explained 59.8% of the variance and the second axis (dim2), 23.6%. Dim1 was positively related to precipitations of the warmest quarter (bio18) and negatively related to mean annual temperature (bio1), mean temperature of the warmest quarter (bio10) and precipitation seasonality (bio15). Dim2 was mostly related to annual precipitation (bio12). Three out of four drought resistance indicators were mostly related to dim1 while Aussenac indicator was related to dim2.

**Figure 3:**
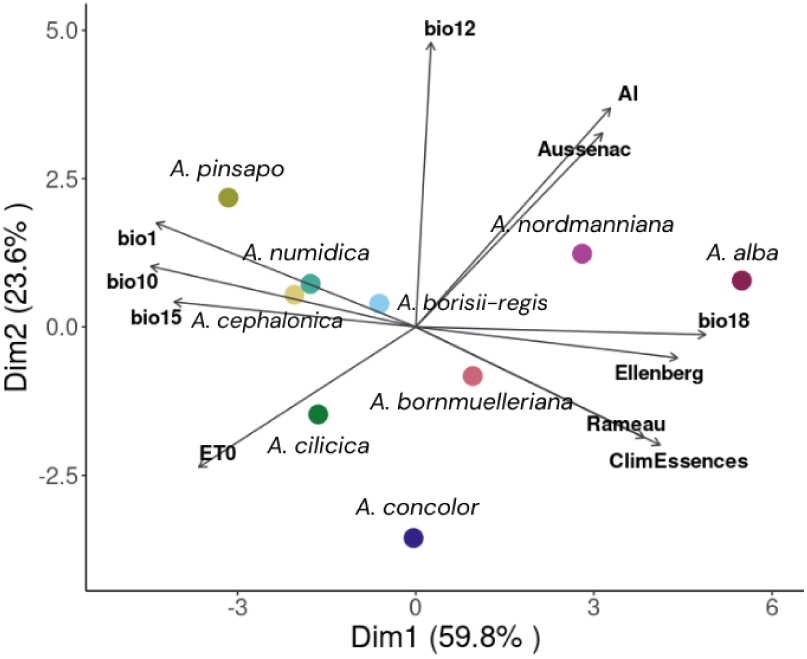
Principal Component Analysis of ”operational” indicators of EuroMediterranean *Abies* species. Bioclimatic variables: bio1 (annual mean temperature), bio10 (mean temperature of warmest quarter), bio12 (annual precipitation), bio15 (pre-cipitation seasonality), bio18 (precipitation of warmest quarter), AI (Aridity Index) and ET0 (potential evapotranspiration). Drought resistance indicators: ClimEssences (mature tree’s resistance to extreme drought), Aussenac (resistance to desiccation of excised branches), Rameau (xericity index), Ellenberg-type indicator. Missing values were set to mean indicator value.

### 3.2. Inter-specific comparison of traits

Some traits presented significant differences between species in their mean, as indicated by the Tukey test presented in figure 4. *A. pinsapo* and *A. numidica* had a *g_min_* 1.8 times lower than *A. concolor* and *A. nordmanniana*. *A. pinsapo* and *A. numidica* had a 1.4-fold higher *THF* and twice as high *SMRI* than *A. concolor*. Mean trait values also differed among species for *SLA*, *HV*, succulence and *SSM* . *A. bornmuelleriana* and *A. nordmanniana* had a significantly higher *SLA* than *A. pinsapo, A. cephalonica* and *A. concolor*. Species differed in their water reserves as *A. cephalonica* and *A. concolor* had a higher succulence than *A. bornmuelleriana, A. nordmanni-ana* and *A. numidica*. *HV* was significantly higher for *A. concolor* than for *A. bornmuelleriana, A. cilicica, A. nordmanniana* and *A. numidica*. *SSM* was significantly higher for *A. nordmanniana* than for *A. bornmuelleriana, A. cephalonica, A. cilicica, A. concolor* and *A. numidica*. *P*_50_ and Π*_T LP_* showed no significant differences between species. Intraspecific variability of the measured traits were not significantly different at the 5% level between species according to Levene’s test (minimal *p_value_* = 0.0508 for *SSM* and *P*_50_ ; *p_value_* = 0.0536 for *g_min_*; *p_value_* = 0.0761 for *SLA* ; for other traits *p_value_ >* 0.3; *data not shown*).

**Figure 4:**
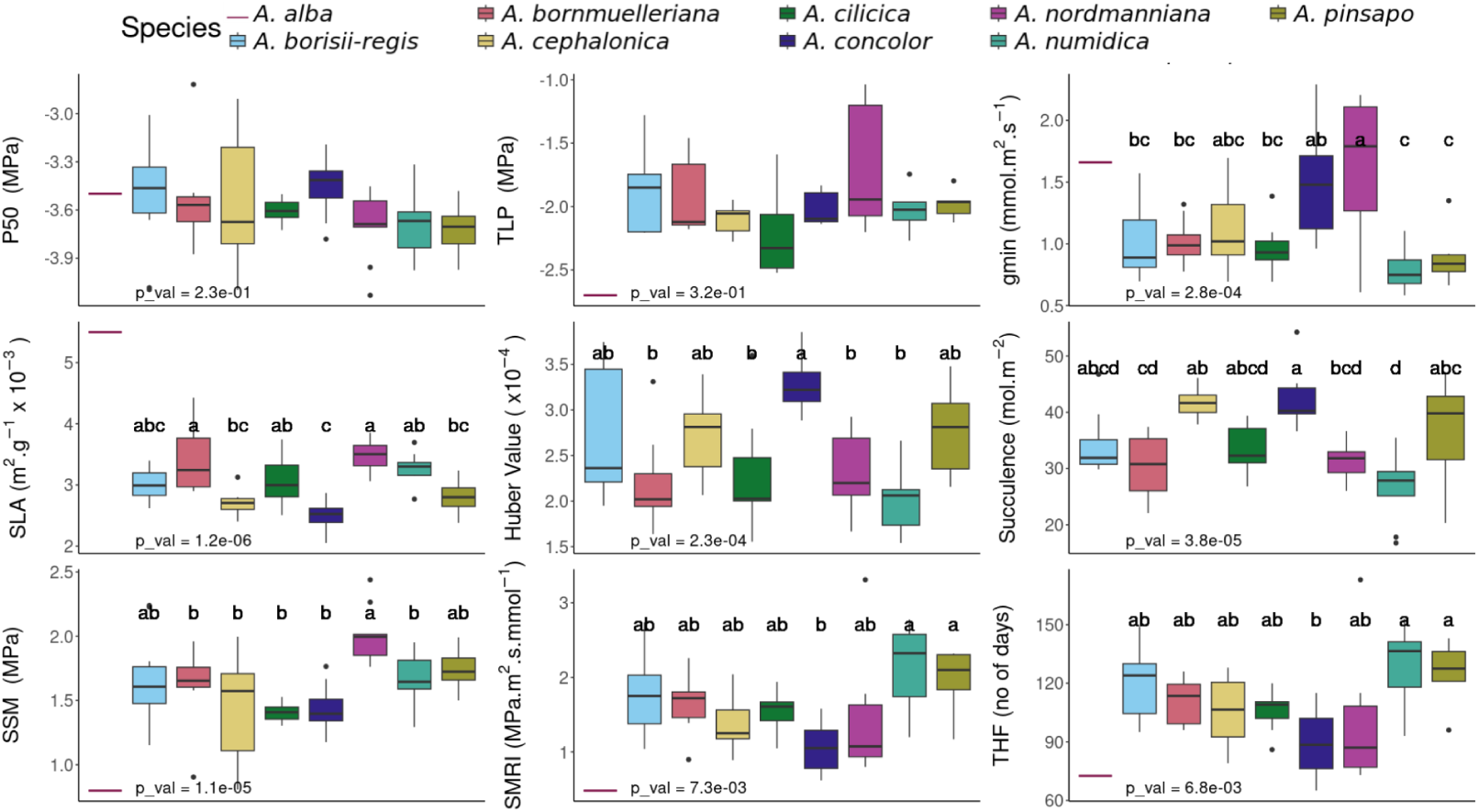
Tukey post-hoc tests for measured hydraulic traits and integrated traits. *P*_50_: water potential at 50% loss of conductivity, Π*_T LP_* : water potential at turgor loss point, *g_min_*: minimal water losses after stomatal closure, *SLA*: Specific Leaf Area, Succulence, Huber Value, *SSM* : Stomatal Safety Margin, *SMRI*: Stomatal Margin Retention Index, *THF* : Time to Hydraulic Failure. *g_min_*, succulence, Huber Value and *SMRI* refer to projected leaf area.

3.3. *Traits explaining vulnerability to hydraulic failure*

Time to Hydraulic Failure was computed and regressed against different traits (figure 5). At the mean species level, with *A. alba*, *g_min_*, *SSM* and *SMRI* are significantly related to *THF* . *g_min_* and *SMRI* showed highest *R*^2^ with *THF* (*R*^2^ = 0.87 and *R*^2^ = 0.93, respectively). At the individual level (figure 5 and supplementary figure C.2), except for Π*_T LP_*, all traits are significantly explicative of *THF* . For *P*_50_, Huber Value and *SSM*, *R*^2^ is below 0.1. For succulence, *R*^2^ = 0.18 and for *g_min_* and *SMRI*, *R*^2^ = 0.76 and *R*^2^ = 0.91 respectively.

**Figure 5:**
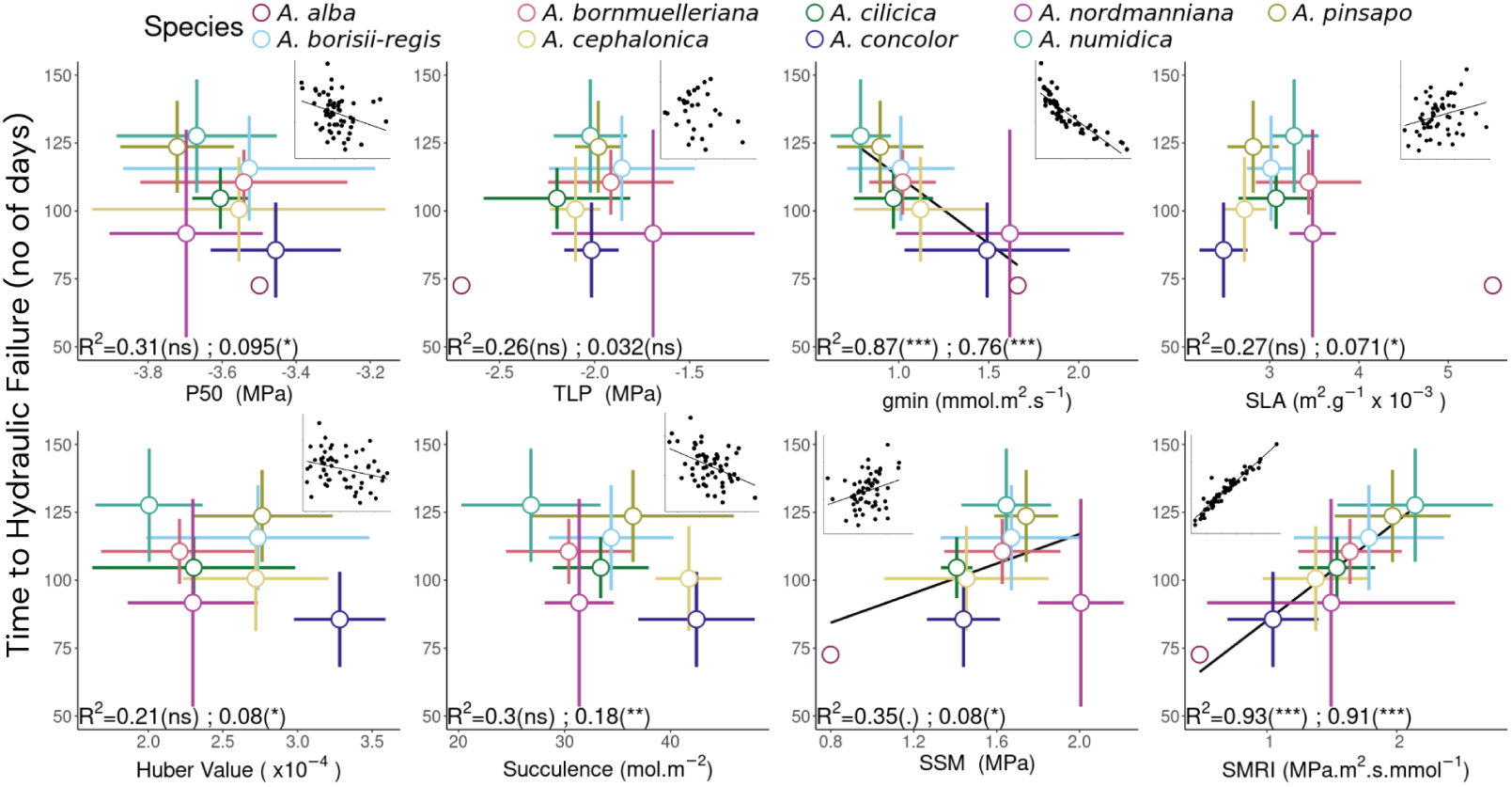
Regression of the Time to Hydraulic Failure on hydraulic traits and two integrated indicators. First regression is performed on the average values of the species, with *Abies alba*. Second regression is performed on individual values. *p_values_* are indicated by the following symbology : ∗∗∗ *<* 0.001 ; ∗∗ *<* 0.05 ; ∗ *<* 0.01, *. <* 0.1. *P*_50_: water potential at 50% loss of conductivity, Π*_T LP_* : water potential at turgor loss point, *g_min_*: minimal water losses after stomatal closure, *SLA*: Specific Leaf Area, Huber Value, succulence, *SSM* : Stomatal Safety Margin, *SMRI*: Stomatal Margin Retention Index.

### 3.4. Relating operational indicators to the vulnerability to hydraulic failure

Time to Hydraulic Failure, computed with SurEau, was regressed on the first dimension of the PCA of operational indicators, showing a significant rela tionship between the two variables (figure 6a): *p_value_* = 0.014 and *R*^2^ = 0.60. *THF* was also regressed against each operational indicators separately (figure 6b), which revealed that the best proxy of *THF* was ClimEssences drought resistance indicator (*p_value_ <* 0.05 and *R*^2^ = 0.75). It is worth mentioning that when *Abies alba* was removed from the dataset, mean annual temperature and ClimEssences were equally good as proxies of *THF* (*p_values_ <* 0.01 and *R*^2^ = 0.58). Correlation of mechanistic traits with operational indicators (supplementary figure C.3) showed that only a few traits related to operational indicators. *P*_50_ was the only trait related to annual precipitation and Π*_T LP_* only related to precipitation of the warmest quarter and PCA-dim.1 when *A. alba* was removed from the dataset.

**Figure 6:**
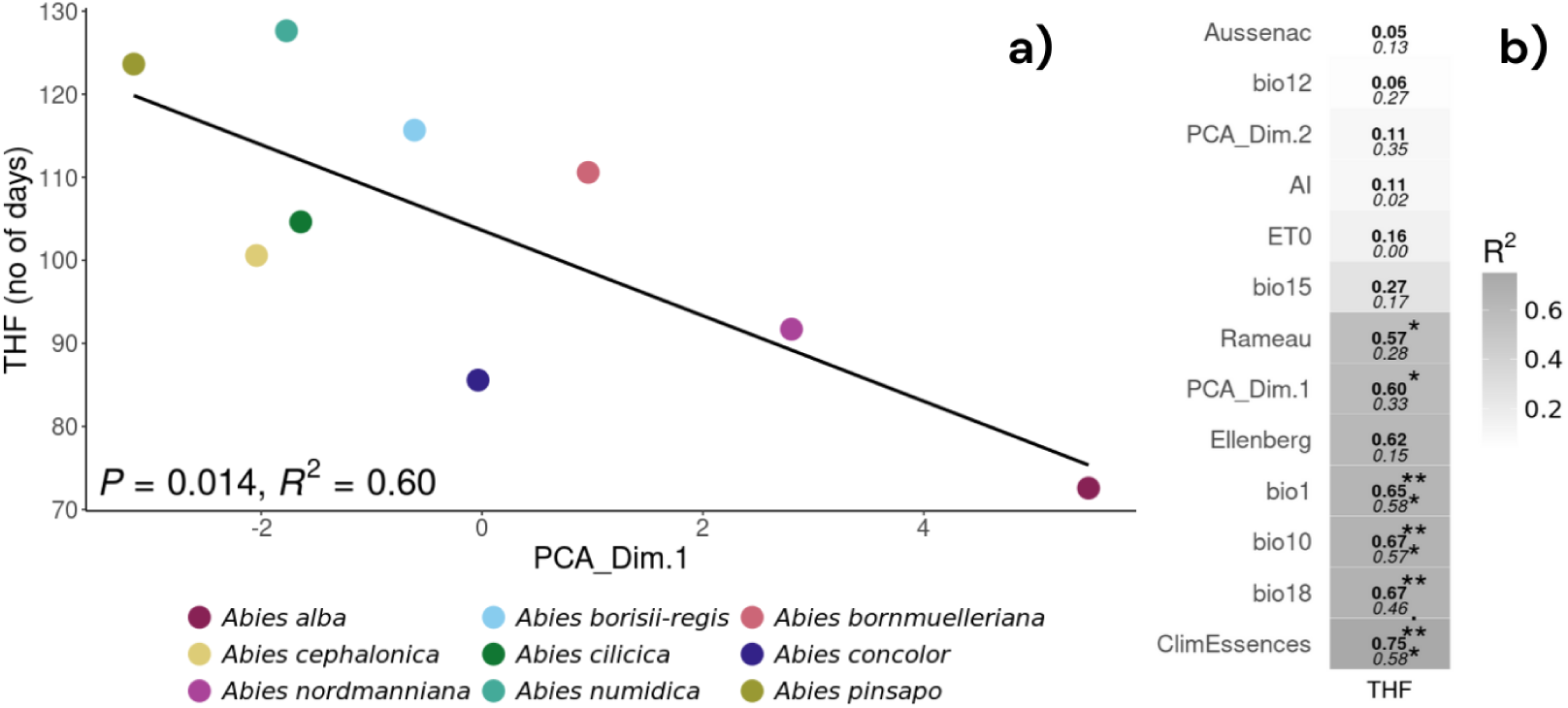
a) Time to Hydraulic Failure (*THF*) regressed on dim1 of Principal Component Analysis of ”operational” indicators. b) *p_value_* and *R*^2^ of *THF* regressed on operational indicators (drought resistance indicators and bioclimatic variables). *p_values_* are indicated by the following symbology : ∗ ∗ ∗ *<* 0.001 ; ∗∗ *<* 0.05 ; ∗ *<* 0.01, *. <* 0.1 For each variable, value with (above) and without (below) *Abies alba* are displayed. Bioclimatic variables: bio1 (annual mean temperature), bio10 (mean temperature of warmest quarter), bio12 (annual precipitation), bio15 (precipitation seasonality), bio18 (precipitation of warmest quarter), AI (Aridity Index) and ET0 (potential evapotranspiration). Drought resistance indicators: ClimEssences (mature tree’s resistance to extreme drought), Aussenac (resistance to desiccation of excised branches), Rameau (xericity index), Ellenberg-type indicator

### 3.5. Climate projection and risk assessment

We highlighted the potential of our ecophysiological approach to estimate the future risk of decline of our different species by computing the risk of hydraulic failure at the end of the century under three climatic scenarios. Figure 7 shows mean species’ hydraulic failure risk in three different Representative Concentration Pathways during the 2070-2100 period. For more than half the years, in all three RCPs, *A. alba* reaches the threshold of 50% loss of conductivity. Similarly, *A. concolor* shows high risk values: the threshold is exceeded in more than one year out of two in the three scenarios. In RCP 2.6, *A. borisii-regis, A. bornmuelleriana, A. numidica* and *A. pinsapo* have a risk of less than 25%. In RCP 4.5, *A. numidica* and *A. pinsapo* have about a one in three chance of reaching 50% loss of conductivity. Nevertheless, in the most extreme scenario, these last two species have a three out of four chance of reaching the threshold.

**Figure 7:**
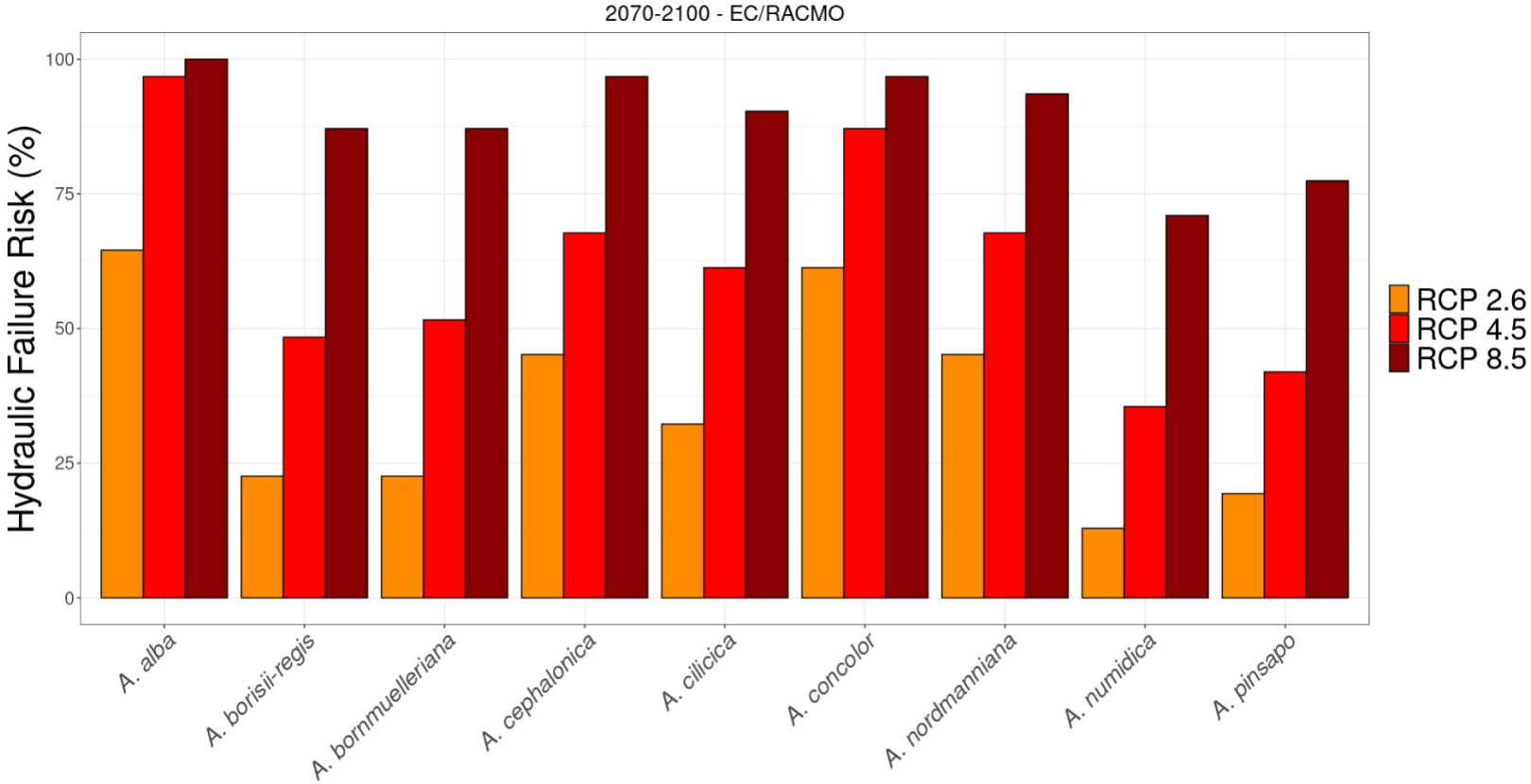
Hydraulic failure risk (probability of exceeding 50% loss of conductivity over the 2070-20100 period) for the different *Abies* species at the experimental plantation of Le Treps, using three Representative Concentration Pathways (RCP) of the EC/RACMO simulation of the DRIAS for the period 2070-2100.

## 4. Discussion

In this study, we assessed whether the use of multiple traits based on species ecophysiology could complement the information provided by ”operational” indicators and help choose species adapted to future climatic conditions. Our work has focused on the Euro-Mediterranean species complex of the *Abies* genus and compared ”operational” indicators to a single indicator derived from ecophysiological modelling: the vulnerability to hydraulic failure (*THF*). Our results shed light on which indicators used operationally for stand adaptation are most associated with *THF* and the prospects and opportunities hydraulic traits offer for improving decision-support tools.

### 4.1. Predicting drought resistance : from single traits, to integrated traits and models

Pioneering literature in ecophysiology has often focused on a single trait to explain species’ resistance to drought. Many studies have focused on resistance to cavitation (*P*_50_) (Choat et al., 2012; Larter et al., 2017; López et al., 2021), turgor loss point (Π*_T LP_*) (Sjöman et al., 2018; Zhu et al., 2018; Hannus et al., 2021) and hydraulic safety margins (*SSM*) (Choat et al., 2012; Martin-StPaul et al., 2017). On a large scale, between biomes and at an inter-specific level, drought resistance has been well explained by species’ resistance to cavitation (Choat et al., 2012; Lens et al., 2016) or by turgor loss point (Bartlett et al., 2012). However, on a smaller scale (*e.g.* at the intra-specific level), resistance to cavitation is not discriminating populations growing in different aridity (Martínez-Vilalta et al., 2009; Lamy et al., 2011; López et al., 2013; Rosas et al., 2019). Moreover, for some genus, not one but several traits correlate with species’ climatic preferences (Larter et al., 2017; Rosas et al., 2019). As underlined by Choat et al. (2018), there is increasing evidence that a single trait does not alone explain drought resistance diversity of closely related species.

In our data, we didn’t find any differences in *P*_50_ within *Abies* species, consistently with previous work from Peguero-Pina et al. (2011). Studying *A. pinsapo* and *A. alba*, they found that both species presented similar resistance to cavitation and that *A. pinsapo*’s higher efficiency of water transport could explain its better drought resistance. However, at the individual level, we found that *P*_50_ was related to *THF* . Regarding turgor loss point (Π*_T LP_*), Petek-Petrik et al. (2023) reported that conifer species with differing drought resistance can show low differentiation for Π*_T LP_* . Again, this is consistent with our findings: this trait does not differentiate our species and does not explain *THF* . By contrast we found that *g_min_* differed between species and was highly explicative of *THF*, which indicates that this trait has a leading role in drought resistance within the *Abies* genus. This is in agreement with recent literature indicating the major role of *g_min_* in resistance to drought (Brodribb et al., 2014; Duursma et al., 2019; Billon et al., 2020; Lanning et al., 2020).

Today, we know that numerous traits influence dehydration dynamics (Blackman et al., 2016; Martin-StPaul et al., 2017; Choat et al., 2018). Literature showed that integrated traits such as *SMRI* (Petek-Petrik et al., 2023) or *THF* (Ruffault et al., 2022) relate well to drought resistance. This is in accordance with our results as we found that these integrated indicators were highly correlated between each other and allowed to differentiate the eight closely related firs. In addition, the relationship between mean bioclimatic variables and *THF* is consistent with previous studies on the response of growth to climate in fir trees. Indeed, by measuring four tree-ring drought indices, George et al. (2015) found that precipitations of wettest quarter and wettest month were best predictors of drought responses in *Abies* species. Our data showed that precipitation of the warmest quarter, mean temperature of warmest quarter and annual mean temperature were well correlated to *THF* . Other studies have found such relationships between traits and bioclimatic variables (Maherali et al., 2004; Choat et al., 2012; Ramírez-Valiente et al., 2020). Species from the *Piceaster* section (*A. pinsapo* and *A. numidica*) have traits conferring more resistance to drought and better drought resistance indicators. These two species come from the southern margins of Mediterranean firs’ range, with climates characterised by long hot and dry periods during the summer. Although in our study, *g_min_* has a leading role on *THF*, other traits were related to *THF* at the individual level. Therefore, integrated indicators based on models make it possible to limit the pitfalls of using a single trait. Whereas several traits are often used as indicators of drought resistance (*P*_50_, Π*_T LP_*, *SSM*), aggregated traits from models allow the integration of the full spectrum of hydraulic and leaf traits depicting diverse strategies of drought resistance (Delzon, 2015).

Our results show that not all operational indicators are good proxies of the vulnerability to hydraulic failure as assessed by integrated physiological indicators (*SMRI*, *THF*). The French indicator from ClimEssences had the highest link to *THF* . This indicator is based on drought-related mortality data and existing literature. For the *Abies* genus, the reference literature used by ClimEssences is the pioneering work in ecophysiology of Aussenac (1980, 2002). Although the indicator proposed by Aussenac (1980) did not relate to *THF*, combining his ecophysiological results to empirical field data provided the best proxy of *THF* : ClimEssences. However, when removing *Abies alba* from the dataset, *p_values_* and *R*^2^ of mean annual temperature and ClimEssences indicator with *THF* were identical, which suggests that the existing relation between *THF* and ClimEssences is mainly due to the strong difference between *Abies alba* and Mediterranean firs. This indicator is not effective for differentiating between closely related species. Aussenac indicator is correlated to the *P*_50_ and annual precipitation (supplementary figure C.3). Aussenac indicator could be related to other mechanisms of drought resistance. Although three ”operational” indicators provided a good estimation of average species vulnerability to hydraulic failure, they do not characterise the diversity of the *Abies* genus.

In a different way, the SurEau model made it possible to take into account intra genus differences with a reduced number of traits and one experimental site. We estimated vulnerability to hydraulic failure associated with a particular site. For the purpose of comparing tree population growing in different conditions of climate and fertility, *THF* could be adjusted by accounting for the evapotranspiration and leaf area index of corresponding sites. The model was also used as a tool for projections in future climatic conditions, beyond empirical knowledge from past and current climate. The ”operational” and the mechanistic approach are complementary for characterising the diversity of drought responses.

#### 4.1.1. Towards the characterisation of intra-specific variability

Bioclimatic values showed that species can occur over a wide range of climatic conditions. For *A. nordmanniana*, precipitation of the warmest quarter ranged from 76 to 375 mm. Species with large range areas (*e.g. A. nordmanniana, A. cephalonica*) could show greater intra-specific trait variability than species with very reduced range areas (*e.g. A. numidica*). Moreover, sampled populations had bioclimatic values outside from their range areas. This underlines that the quality of presence/absence data is another limitation to the accuracy of distribution models (Lobo, 2008). The small size of our sampling (ten individuals per species) didn’t allow us to detect differences of intra-specific variability between species. However, Levene’s test *p_values_* for *SSM*, *P*_50_ and *g_min_* were very close to the 5% threshold indicating that further studies, with larger samples, are required to confirm that intra-specific variances are different. Indeed, species such as *A. nordmanniana* do seem to have a high variance relatively to other species regarding *g_min_*. IPCC defines the vulnerability as ”the sensitivity or susceptibility to harm and lack of capacity to cope and adapt” (IPCC, 2018). Intra-specific variability being one of the main drivers of adaptation (Śanchez-Gómez and Aranda, 2024), our study calls for a more in-depth investigation of the intraspecific variability of vulnerability to hydraulic failure.

#### 4.1.2. Consequences for fir ecology and management

Our results indicate that species from the *Piceaster* section (*A. pinsapo, A. numidica*) are less vulnerable to hydraulic failure than others. The ability to withstand long summer droughts has been reported for *A. pinsapo* (Linares, 2011). Its drought resistance was linked by Peguero-Pina et al. (2011) to the species’ wider tracheids. With wider tracheids, *A. pinsapo* is more efficient at transporting water through the xylem, enabling good water supply in conditions of high atmospheric evaporative demand, typical of Mediterranean summers (Peguero-Pina et al., 2011). However, this makes *A. pinsapo* more sensitive to freeze-thaw induced cavitation (Pittermann and Sperry, 2006). Indeed, Aussenac (2002) already reported the risk of late frost damage for other Mediterranean *Abies* species (*A. cephalonica* and *A. cilicica*). We also noted that the largest trees had the highest *P*_50_ and were therefore less resistant to cavitation (data not shown). Phenology and growth are other aspects of trees life-history traits that require further investigation for adapted management under climate change. As underlined by Benito-Garźon et al. (2013), major planting/enrichment failures can occur when focusing on a single aspect of species’ climatic suitability. Our results show that traits can be used at species level to estimate vulnerability to hydraulic failure, an indicator of drought resistance, in a way that is globally consistent with drought resistance indicators. Additionally, by integrating traits into a model, they can also be used to assess the behaviour of species in future climates. These trait-based models could thus complement Species Distribution Models currently in use (Alba-Śanchez et al., 2010; López-Tirado et al., 2023), which often operate at the species scale and are based on a subsample of the potential distribution of species. Moreover, with SurEau, we could characterise intra-specific variability of the vulnerability to hydraulic failure, although the approach needs to be validated for the intra-specific level.

## 5. Conclusion

Indicators used operationally by foresters (bioclimatic variables, Ellenbergtype indicators, empirical indicators) are mere proxies of the vulnerability to hydraulic failure. As they summarise a species to a single number, based on their observed response to past and current climate, they do not differentiate closely related species and are not available for all species. For instance, the expert-based ClimEssences indicator informs on the vulnerability to hydraulic failure of *Abies alba* compared to other species but is less accurate to compare among Mediterranean species. As a complement to these indicators, hydraulic traits are great tools for assessing vulnerability and risk in future climates and for subsequent improvement of decision-making tools. Our study showed that in the Euro-Mediterranean fir species complex, vulnerability to hydraulic failure is mainly driven by minimal water losses after stomatal closure, *g_min_*. *Abies pinsapo* and *Abies numidica*, only representatives of the *Piceaster* section, showed the least vulnerability to hydraulic failure. Their natural environment is characterised by long summer droughts and high mean annual temperature. This suggest that one of the mechanisms of diversification of the *Abies* genus in the Mediterranean basin is adaptation to long summer water shortage. Our study focused on one or two provenances per species. A thorough investigation of intra-specific variability needs to be made to assess the possible evolution of *Abies* species in regard of climate change. In the end, integrated indicators of drought resistance such as Stomatal Margin Retention Index (*SMRI*) or Time to Hydraulic Failure (*THF*), estimated with the SurEau model, are good predictors of drought resistance under various scenarios of future climate. These indicators provide complementary information to operational indicators. There wider use in the selection of future forest species or provenances could bring interesting prospects to forest management under climate change.

## List of figures

- Figure 1: Map of the origin of sampled fir trees.
- Figure 2: Conceptual framework of the modelling with the SurEau hydraulic model.
- Figure 3: Principal component analysis of ”operational” indicators.
- Figure 4: Tukey post-hoc test for measured hydraulic traits and integrated traits.
- Figure 5: Regression of Time to Hydraulic Failure on hydraulic traits and two integrated indicators.
- Figure 6: Time to Hydraulic Failure regressed on the first dimension of the PCA of operational indicators.
- Figure 7: Climatic projections of the hydraulic failure risk.

## List of supplementary data

- Table 1: Abies species sampled in the two experimental setups.
- Table 2: Source of trait values for Abies alba
- Table 3: First setup composition.
- Table 4: Second setup composition.
- Table 5: Bioclimatic variables of Abies species.
- Table 6: Drought resistance indicators of Abies species.
- Figure 1: QQPlot for Anova analysis.
- Figure 2: Individual regression of time to hydraulic failure on traits.
- Figure 3: Correlations between ecophysiological traits and operational indicators.

## Glossary

*HV* Huber Value. 11, 16, 17

*IPCC* Intergovernmental Panel on Climate Change. 13

*P*_50_ Water potential at 50% loss of conductivity. 4, 9, 12, 13, 17–19, 21–24, 39, 40, 45, 46

*RWC_T LP_* Relative Water Content at Turgor Loss Point. 9

*S* Slope of the relationship between water potential and water content when water potential is equal to *P*_50_ . 9, 13, 40

*SLA* Specific Leaf Area. 11, 16–18, 45, 46

*SMRI* Stomatal Margin Retention Index: indicator of the time required to cross the stomatal safety margin. 4, 10, 16–18, 22, 25, 45, 46

*SSM* Stomatal Safety Margin: 0.95 × *ψ_tlp_* − Ψ_50_. 4, 10, 16–18, 21–23, 45, 46

*THF* Time to Hydraulic Failure. 1, 5, 13, 14, 16–19, 21–23, 25, 46

Π*_T LP_* Water potential at Turgor Loss Point. 4, 9, 17–19, 21, 22, 40, 45, 46 Ψ*_gs_*_12_

Water potential in leaves for 12% stomatal closure. 12, 13

Ψ*_gs_*_88_ Water potential in leaves for 88% stomatal closure. 12, 13

*ɛ* Elasticity modulus. 9, 12, 13, 40

*π*_0_ Osmotic potential at full turgor. 9, 12, 13, 40

*a_f_* Apoplasmic fraction. 9, 12, 13, 40

*g_min_* Minimal water losses after stomatal closure. 4, 9, 10, 12, 13, 16–18, 21–25, 40, 45, 46

*gs_max_* Maximum stomatal conductance. 12, 13

## References

1. Adams, H.D., Zeppel, M.J.B., Anderegg, W.R.L., Hartmann, H., Landhäusser, S.M., Tissue, D.T., Huxman, T.E., Hudson, P.J., Franz, T.E., Allen, C.D., Anderegg, L.D.L., Barron-Gafford, G.A., Beerling, D.J., Breshears, D.D., Brodribb, T.J., Bugmann, H., Cobb, R.C., Collins, A.D., Dickman, L.T., Duan, H., Ewers, B.E., Galiano, L., Galvez, D.A., Garcia-Forner, N., Gaylord, M.L., Germino, M.J., Gessler, A., Hacke, U.G., Hakamada, R., Hector, A., Jenkins, M.W., Kane, J.M., Kolb, T.E., Law, D.J., Lewis, J.D., Limousin, J.M., Love, D.M., Macalady, A.K., MartínezVilalta, J., Mencuccini, M., Mitchell, P.J., Muss, J.D., O’Brien, M.J., O’Grady, A.P., Pangle, R.E., Pinkard, E.A., Piper, F.I., Plaut, J.A., Pockman, W.T., Quirk, J., Reinhardt, K., Ripullone, F., Ryan, M.G., Sala, A., Sevanto, S., Sperry, J.S., Vargas, R., Vennetier, M., Way, D.A., Xu, C., Yepez, E.A., McDowell, N.G., 2017. A multi-species synthesis of physiological mechanisms in drought-induced tree mortality. Nat. Ecol. Evol. 1, 1285–1291. 10.1038/s41559-017-0248-x.

2. Aitken, S.N., Bemmels, J.B., 2016. Time to get moving: assisted gene flow of forest trees. Evol. Appl. 9, 271–290. 10.1111/eva.12293.

3. Aitken, S.N., Whitlock, M.C., 2013. Assisted gene flow to facilitate local adaptation to climate change. Annu. Rev. Ecol. Evol. Syst. 44, 367–388. 10.1146/annurev-ecolsys-110512-135747.

4. Alba-Sánchez, F., Ĺopez-Sáez, J.A., Pando, B.B.d., Linares, J.C., Nieto-Lugilde, D., López-Merino, L., 2010. Past and present potential distribution of the Iberian Abies species: a phytogeographic approach using fossil pollen data and species distribution models. Divers. distrib. 16, 214–228. 10.1111/j.1472-4642.2010.00636.x.

5. Allen, C.D., Breshears, D.D., McDowell, N.G., 2015. On underestimation of global vulnerability to tree mortality and forest die-off from hotter drought in the anthropocene. Ecosphere 6, 129. 10.1890/ES15-00203.1.

6. Aussenac, G., 1980. Comportement hydrique de rameaux exciśes de quelques espèces de sapins et de pins noirs en phase de dessiccation. Ann. For. Sci. 37, 201–215. 10.1051/forest:19800303.

7. Aussenac, G., 2002. Ecology and ecophysiology of circum-mediterranean firs in the context of climate change. Ann. For. Sci. 59, 823–832. 10.1051/forest:2002080.

8. Ay, J.S., Guillemot, J., Martin-StPaul, N., Doyen, L., Leadley, P., 2017. The economics of land use reveals a selection bias in tree species distribution models. Global Ecol. Biogeogr. 26, 65–77. 10.1111/geb.12514.

9. Balao, F., Lorenzo, M.T., Sánchez-Robles, J.M., Paun, O., GarćıaCastaño, J.L., Terrab, A., 2020. Early diversification and permeable species boundaries in the mediterranean firs. Ann. Bot. 125, 495–507. 10.1093/aob/mcz186.

10. Bartlett, M.K., Scoffoni, C., Sack, L., 2012. The determinants of leaf turgor loss point and prediction of drought tolerance of species and biomes: a global meta-analysis. Ecol. Lett. 15, 393–405. 10.1111/j.1461-0248.2012.01751.x.

11. Benito-Garzón, M., Ha-Duong, M., Frascaria-Lacoste, N., FerńandezManjarŕes, J.F., 2013. Extreme climate variability should be considered in forestry assisted migration. BioScience 63, 317. 10.1525/bio.2013.63.5.20.

12. Billon, L.M., Blackman, C.J., Cochard, H., Badel, E., Hitmi, A., Cartailler, J., Souchal, R., Torres-Ruiz, J.M., 2020. The DroughtBox: A new tool for phenotyping residual branch conductance and its temperature dependence during drought. Plant Cell Environ. 43, 1584–1594. 10.1111/pce.13750.

13. Blackman, C.J., Pfautsch, S., Choat, B., Delzon, S., Gleason, S.M., Duursma, R.A., 2016. Toward an index of desiccation time to tree mortality under drought. Plant Cell Environ. 39, 2342–2345. 10.1111/pce.12758.

14. Brodribb, T.J., Holbrook, N.M., 2003. Stomatal closure during leaf dehydration, correlation with other leaf physiological traits. Plant Physiol. 132, 2166–2173. 10.1104/pp.103.023879.

15. Brodribb, T.J., McAdam, S.A.M., Jordan, G.J., Martins, S.C.V., 2014. Conifer species adapt to low-rainfall climates by following one of two divergent pathways. Proc. Natl. Acad. Sci. U.S.A. 111, 14489–14493. 10.1073/pnas.1407930111.

16. Chakraborty, D., Ciceu, A., Ballian, D., Benito Garzón, M., Bolte, A., Bozic, G., Buchacher, R., C^̌^epl, J., Cremer, E., Ducousso, A., Gaviria, J., George, J.P., Hardtke, A., Ivankovic, M., Klisz, M., Kowalczyk, J., Kremer, A., Lstiburek, M., Longauer, R., Mihai, G., Nagy, L., Petkova, K., Popov, E., Schirmer, R., Skrøppa, T., Solvin, T.M., Steffenrem, A., Stejskal, J., Stojnic, S., Volmer, K., Schueler, S., 2024. Assisted tree migration can preserve the european forest carbon sink under climate change. Nat. Clim. Change. 10.1038/s41558-024-02080-5.

17. Chave, J., Coomes, D., Jansen, S., Lewis, S.L., Swenson, N.G., Zanne, A.E., 2009. Towards a worldwide wood economics spectrum. Ecol. Lett. 12, 351–366. 10.1111/j.1461-0248.2009.01285.x.

18. Choat, B., Brodribb, T.J., Brodersen, C.R., Duursma, R.A., Ĺopez, R., Medlyn, B.E., 2018. Triggers of tree mortality under drought. Nature 558, 531–539. 10.1038/s41586-018-0240-x.

19. Choat, B., Jansen, S., Brodribb, T.J., Cochard, H., Delzon, S., Bhaskar, R., Bucci, S.J., Feild, T.S., Gleason, S.M., Hacke, U.G., Jacobsen, A.L., Lens, F., Maherali, H., Martínez-Vilalta, J., Mayr, S., Mencuccini, M., Mitchell, P.J., Nardini, A., Pittermann, J., Pratt, R.B., Sperry, J.S., Westoby, M., Wright, I.J., Zanne, A.E., 2012. Global convergence in the vulnerability of forests to drought. Nature 491, 752–755. 10.1038/nature11688.

20. Cochard, H., 2002. A technique for measuring xylem hydraulic conductance under high negative pressures. Plant Cell Environ. 25, 815–819. 10.1046/j.1365-3040.2002.00863.x.

21. Cochard, H., Pimont, F., Ruffault, J., Martin-StPaul, N., 2021. SurEau: a mechanistic model of plant water relations under extreme drought. Ann. For. Sci. 78, 55. 10.1007/s13595-021-01067-y.

22. Delzon, S., 2015. New insight into leaf drought tolerance. Funct. Ecol. 29, 1247–1249. 10.1111/1365-2435.12500.

23. Duursma, R.A., Blackman, C.J., Lopéz, R., Martin-StPaul, N.K., Cochard, H., Medlyn, B.E., 2019. On the minimum leaf conductance: its role in models of plant water use, and ecological and environmental controls. New Phytol. 221, 693–705. 10.1111/nph.15395.

24. Ellenberg, H., Leuschner, C., 2010. Vegetation Mitteleuropas mit den Alpen. 6 .Aufl. Ulmer Verlag.

25. Fady, B., Scotti-Saintagne, C., Cano, M., Riou, A., 2023. Fifty years (1965-2023) of phenotypic trait data for Mediterranean Abies species from Turkey and the Caucasus in French common gardens. Recherche Data Gouv 10.57745/WMRL9Z.

26. Fady, B., Vauthier, D., Rei, F., Scotti-Saintagne, C., 2024. Phenotypic trait data (1972-2023) for the Mediterranean firs Abies cephalonica and A. borisii-regis from eight French common gardens. Recherche Data Gouv 10.57745/XEYITB.

27. Fick, S., Hijmans, R., 2017. Worldclim 2: new 1km spatial resolution climate surfaces for global land areas. Int. J. Climatol. 37, 4302–4315. 10.1002/joc.5086.

28. Fuchs, S., Schuldt, B., Leuschner, C., 2021. Identification of drought-tolerant tree species through climate sensitivity analysis of radial growth in central european mixed broadleaf forests. For. Ecol. Manag. 494, 119287. 10.1016/j.foreco.2021.119287.

29. Gaitán-Espitia, J.D., Hobday, A.J., 2021. Evolutionary principles and genetic considerations for guiding conservation interventions under climate change. Glob. Change Biol. 27, 475–488. 10.1111/gcb.15359.

30. George, J.P., Schueler, S., Karanitsch-Ackerl, S., Mayer, K., Klumpp, R.T., Grabner, M., 2015. Inter- and intra-specific variation in drought sensitivity in Abies spec. and its relation to wood density and growth traits. Agric. For. Meteorol. 214–215, 430–443. 10.1016/j.agrformet.2015.08.268.

31. Hannus, S., Hirons, A., Baxter, T., McAllister, H.A., Wiström, B., Sjöman, H., 2021. Intraspecific drought tolerance of betula pendula genotypes: an evaluation using leaf turgor loss in a botanical collection. Trees 35, 569–581. 10.1007/s00468-020-02059-7.

32. Hinckley, T.M., Duhme, F., Hinckley, A.R., Richter, H., 1980. Water relations of drought hardy shrubs: osmotic potential and stomatal reactivity. Plant Cell Environ. 3, 131–140. 10.1111/1365-3040.ep11580919.

33. IPCC, 2014. Climate change 2014: Synthesis report. Intergovernmental Panel on Climate Change, Geneva, Switzerland, 151.

34. IPCC, 2018. Global warming of 1.5°c. Intergovernmental Panel on Climate Change, Geneva, Switzerland. https://www.ipcc.ch/sr15/chapter/spm. Viewed 16 Jan 2025.

35. Jacob, D., Petersen, J., Eggert, B., Alias, A., Christensen, O.B., Bouwer, L.M., Braun, A., Colette, A., Déqúe, M., Georgievski, G., Georgopoulou, E., Gobiet, A., Menut, L., Nikulin, G., Haensler, A., Hempelmann, N., Jones, C., Keuler, K., Kovats, S., Kröner, N., Kotlarski, S., Kriegsmann, A., Martin, E., van Meijgaard, E., Moseley, C., Pfeifer, S., Preuschmann, S., Radermacher, C., Radtke, K., Rechid, D., Rounsevell, M., Samuelsson, P., Somot, S., Soussana, J.F., Teichmann, C., Valentini, R., Vautard, R., Weber, B., Yiou, P., 2014. EURO-CORDEX: new high-resolution climate change projections for European impact research. Reg. Environ. Change 14, 563–578. 10.1007/s10113-013-0499-2.

36. Jiménez-Valverde, A., Lobo, J.M., Hortal, J., 2008. Not as good as they seem: the importance of concepts in species distribution modelling. Divers. Distrib. 14, 885–890. 10.1111/j.1472-4642.2008.00496.x.

37. Jones, H., 1992. Plants and microclimate: a quantitative approach to environmental plant physiology. Cambridge University Press, Cambridge. 10.1017/CBO9780511845727.

38. Jourńe, V., Barnagaud, J., Bernard, C., Crochet, P., Morin, X., 2020. Correlative climatic niche models predict real and virtual species distributions equally well. Ecology 101. 10.1002/ecy.2912.

39. Klein, T., 2014. The variability of stomatal sensitivity to leaf water potential across tree species indicates a continuum between isohydric and anisohydric behaviours. Funct. Ecol. 28, 1313–1320. 10.1111/1365-2435.12289.

40. Korosuo, A., Pilli, R., Abad Viñas, R., Blujdea, V.N.B., Colditz, R.R., Fiorese, G., Rossi, S., Vizzarri, M., Grassi, G., 2023. The role of forests in the EU climate policy: are we on the right track? Carbon Balance Manag. 18, 15. 10.1186/s13021-023-00234-0.

41. Krajmerová, D., Paule, L., Zhelev, P., Voleková, M., Evtimov, I., Gagov, V., Gömöry, D., 2016. Natural hybridization in eastern-mediterranean firs: The case of *Abies borisii-regis*. Plant Biosyst. 150, 1189–1199. 10.1080/11263504.2015.1011723.

42. Kramer, P., 1983. Water relations of plants. Academic Press, New York.

43. Kremer, A., Ronce, O., Robledo-Arnuncio, J.J., Guillaume, F., Bohrer, G., Nathan, R., Bridle, J.R., Gomulkiewicz, R., Klein, E.K., Ritland, K., Kuparinen, A., Gerber, S., Schueler, S., 2012. Long-distance gene flow and adaptation of forest trees to rapid climate change. Ecol. Lett. 15, 378–392. 10.1111/j.1461-0248.2012.01746.x.

44. Kunert, N., Tomaskova, I., 2020. Leaf turgor loss point at full hydration for 41 native and introduced tree and shrub species from Central Europe. J. Plant Ecol. 13, 754–756. 10.1093/jpe/rtaa059.

45. Kurz, M., Kölz, A., Gorges, J., Pablo Carmona, B., Brang, P., Vitasse, Y., Kohler, M., Rezzonico, F., Smits, T.H.M., Bauhus, J., Rudow, A., Kim Hansen, O., Vatanparast, M., Sevik, H., Zhelev, P., Gömöry, D., Paule, L., Sperisen, C., Csilĺery, K., 2023. Tracing the origin of Oriental beech stands across Western Europe and reporting hybridization with European beech – Implications for assisted gene flow. For. Ecol. Manag. 531. 10.1016/j.foreco.2023.120801.

46. Lamy, J.B., Bouffier, L., Burlett, R., Plomion, C., Cochard, H., Delzon, S., 2011. Uniform selection as a primary force reducing population genetic differentiation of cavitation resistance across a species range. PLOS ONE 6. 10.1371/journal.pone.0023476.

47. Lanning, M., Wang, L., Novick, K.A., 2020. The importance of cuticular permeance in assessing plant water–use strategies. Tree Physiol. 40, 425– 432. 10.1093/treephys/tpaa020.

48. Larter, M., Pfautsch, S., Domec, J., Trueba, S., Nagalingum, N., Delzon, S., 2017. Aridity drove the evolution of extreme embolism resistance and the radiation of conifer genus *Callitris*. New Phytol. 215, 97–112. 10.1111/nph.14545.

49. Lens, F., Picon-Cochard, C., Delmas, C.E., Signarbieux, C., Buttler, A., Cochard, H., Jansen, S., Chauvin, T., Chacon Doria, L., Del Arco, M., Delzon, S., 2016. Herbaceous angiosperms are not more vulnerable to drought-induced embolism than angiosperm trees. Plant Physiol. 172, 661– 667. 10.1104/pp.16.00829.

50. Limousin, J.M., Roussel, A., Rodŕıguez-Calcerrada, J., Torres-Ruiz, J.M., Moreno, M., Garcia de Jalon, L., Ourcival, J.M., Simioni, G., Cochard, H., Martin-StPaul, N., 2022. Drought acclimation of quercus ilex leaves improves tolerance to moderate drought but not resistance to severe water stress. Plant Cell Environ. 45, 1967–1984. 10.1111/pce.14326.

51. Linares, J.C., 2011. Biogeography and evolution of abies (pinaceae) in the mediterranean basin: the roles of long-term climatic change and glacial refugia: Biogeography and evolution of the circum-mediterranean firs. J. Biogeogr. 38, 619–630. 10.1111/j.1365-2699.2010.02458.x.

52. Lloret, F., Keeling, E.G., Sala, A., 2011. Components of tree resilience: effects of successive low-growth episodes in old ponderosa pine forests. Oikos 120, 1909–1920. 10.1111/j.1600-0706.2011.19372.x.

53. Lobo, J.M., 2008. More complex distribution models or more representative data? Biodivers. Inform. 5. 10.17161/bi.v5i0.40.

54. Lopez-Iglesias, B., Villar, R., Poorter, L., 2014. Functional traits predict drought performance and distribution of Mediterranean woody species. Acta Oecol. 56, 10–18. 10.1016/j.actao.2014.01.003.

55. López, R., Cano, F.J., Martin-StPaul, N.K., Cochard, H., Choat, B., 2021. Coordination of stem and leaf traits define different strategies to regulate water loss and tolerance ranges to aridity. New Phytol. 230, 497–509. 10.1111/nph.17185.

56. López, R., López De Heredia, U., Collada, C., Cano, F.J., Emerson, B.C., Cochard, H., Gil, L., 2013. Vulnerability to cavitation, hydraulic efficiency, growth and survival in an insular pine (Pinus canariensis). Ann. Bot. 111, 1167–1179. 10.1093/aob/mct084.

57. López-Tirado, J., Moreno-Garćıa, M., Romera-Romera, D., Zarco, V., Hidalgo, P.J., 2023. Forecasting the circum-Mediterranean firs (Abies spp., Pinaceae) distribution: an assessment of a threatened conifers’ group facing climate change in the twenty-first century. New For. 55, 143–156. 10.1007/s11056-023-09972-y.

58. Maherali, H., Pockman, W.T., Jackson, R.B., 2004. Adaptive variation in the vulnerability of woody plants to xylem cavitation. Ecology 85, 2184–2199. 10.1890/02-0538.

59. Martin, R.A., da Silva, C.R.B., Moore, M.P., Diamond, S.E., 2023. When will a changing climate outpace adaptive evolution? WIREs Clim. Change 14. 10.1002/wcc.852.

60. Martin-StPaul, N., Delzon, S., Cochard, H., 2017. Plant resistance to drought depends on timely stomatal closure. Ecol. Lett. 20, 1437–1447. 10.1111/ele.12851.

61. Martínez-Vilalta, J., Cochard, H., Mencuccini, M., Sterck, F., Herrero, A., Korhonen, J.F.J., Llorens, P., Nikinmaa, E., Nolè, A., Poyatos, R., Ripullone, F., Sass-Klaassen, U., Zweifel, R., 2009. Hydraulic adjustment of Scots pine across Europe. New Phytol. 184, 353–364. 10.1111/j.1469-8137.2009.02954.x.

62. Mas, E., Cochard, H., Deluigi, J., Didion-Gency, M., Martin-StPaul, N., Morcillo, L., Valladares, F., Vilagrosa, A., Grossiord, C., 2024. Interactions between beech and oak seedlings can modify the effects of hotter droughts and the onset of hydraulic failure. New Phytol. 241, 1021–1034. 10.1111/nph.19358.

63. Matías, L., Quero, J.L., Zamora, R., Castro, J., 2012. Evidence for plant traits driving specific drought resistance. a community field experiment. Environ. Exp. Bot. 81, 55–61. 10.1016/j.envexpbot.2012.03.002.

64. Moreno, M., Simioni, G., Cochard, H., Doussan, C., Guillemot, J., Decarsin, R., Fernandez-Conradi, P., Dupuy, J.L., Trueba, S., Pimont, F., Ruffault, J., Jean, F., Marloie, O., Martin-StPaul, N.K., 2024. Isohydricity and hydraulic isolation explain reduced hydraulic failure risk in an experimental tree species mixture. Plant Physiol. 195, 2668–2682. 10.1093/plphys/kiae239.

65. Niinemets, U., Valladares, F., 2006. Tolerance to shade, drought, and waterlogging of temperate northern hemisphere trees and shrubs. Ecol. Monogr. 76, 521–547.

66. O’Brien, M.J., Engelbrecht, B.M.J., Joswig, J., Pereyra, G., Schuldt, B., Jansen, S., Kattge, J., Landhäusser, S.M., Levick, S.R., Preisler, Y., Väanänen, P., Macinnis-Ng, C., 2017. A synthesis of tree functional traits related to drought-induced mortality in forests across climatic zones. J. Appl. Ecol 54, 1669–1686. 10.1111/1365-2664.12874.

67. Peguero-Pina, J.J., Sancho-Knapik, D., Cochard, H., Barredo, G., Villarroya, D., Gil-Pelegrin, E., 2011. Hydraulic traits are associated with the distribution range of two closely related Mediterranean firs, Abies alba Mill. and Abies pinsapo Boiss. Tree Physiol. 31, 1067–1075. 10.1093/treephys/tpr092.

68. Petek-Petrik, A., Petŕık, P., Lamarque, L.J., Cochard, H., Burlett, R., Delzon, S., 2023. Drought survival in conifer species is related to the time required to cross the stomatal safety margin. J. Exp. Bot. 74, 6847–6859. 10.1093/jxb/erad352.

69. Peterson St-Laurent, G., Hagerman, S., Kozak, R., 2018. What risks matter? Public views about assisted migration and other climate-adaptive reforestation strategies. Clim. Change 151, 573–587. 10.1007/s10584-018-2310-3.

70. Piedallu, C., Dallery, D., Bresson, C., Legay, M., Gégout, J.C., Pierrat, R., 2023. Spatial vulnerability assessment of silver fir and norway spruce dieback driven by climate warming. Landsc. Ecol. 38, 341–361. 10.1007/s10980-022-01570-1.

71. Pittermann, J., Sperry, J.S., 2006. Analysis of freeze-thaw embolism in conifers. the interaction between cavitation pressure and tracheid size. Plant Physiol. 140, 374–382. 10.1104/pp.105.067900.

72. Quigley, K.M., Bay, L.K., van Oppen, M.J.H., 2019. The active spread of adaptive variation for reef resilience. Ecol. Evol. 9, 11122–11135. 10.1002/ece3.5616.

73. Radchuk, V., Reed, T., Teplitsky, C., van de Pol, M., Charmantier, A., Hassall, C., Adamík, P., Adriaensen, F., Ahola, M.P., Arcese, P., Miguel Aviĺes, J., Balbontin, J., Berg, K.S., Borras, A., Burthe, S., Clobert, J., Dehnhard, N., de Lope, F., Dhondt, A.A., Dingemanse, N.J., Doi, H., Eeva, T., Fickel, J., Filella, I., Fossøy, F., Goodenough, A.E., Hall, S.J.G., Hansson, B., Harris, M., Hasselquist, D., Hickler, T., Joshi, J., Kharouba, H., Martínez, J.G., Mihoub, J.B., Mills, J.A., Molina-Morales, M., Moksnes, A., Ozgul, A., Parejo, D., Pilard, P., Poisbleau, M., Rousset, F., Rödel, M.O., Scott, D., Senar, J.C., Stefanescu, C., Stokke, B.G., Kusano, T., Tarka, M., Tarwater, C.E., Thonicke, K., Thorley, J., Wilting, A., Tryjanowski, P., Merila, J., Sheldon, B.C., Pape Møller, A., Matthysen, E., Janzen, F., Dobson, F.S., Visser, M.E., Beissinger, S.R., Courtiol, A., Kramer-Schadt, S., 2019. Adaptive responses of animals to climate change are most likely insufficient. Nat. Commun. 10, 3109. 10.1038/s41467-019-10924-4.

74. Rameau, J.C., Mansion, D., Dumé, G., Gauberville, C., 2016. Flore Forestìere Fraņcaise volume 3. CNPF, Institut pour le développement Forestier.

75. Ramírez-Valiente, J.A., López, R., Hipp, A.L., Aranda, I., 2020. Correlated evolution of morphology, gas exchange, growth rates and hydraulics as a response to precipitation and temperature regimes in oaks (*Quercus*). New Phytol. 227, 794–809. 10.1111/nph.16320.

76. RMT AFORCE, 2021. Fiches espèces du ŕeseau mixte technologique d’accompagnement des forestiers dans l’adaptation des for^ets aux changements climatiques. https://climessences.fr/fiches-especes/fiches-especes. Accessed: 2024-05-24.

77. Rosas, T., Mencuccini, M., Barba, J., Cochard, H., Saura-Mas, S., MartínezVilalta, J., 2019. Adjustments and coordination of hydraulic, leaf and stem traits along a water availability gradient. New Phytol. 223, 632–646. 10.1111/nph.15684.

78. Ruffault, J., Pimont, F., Cochard, H., Dupuy, J.L., Martin-StPaul, N., 2022. SurEau-ecos v2.0: a trait-based plant hydraulics model for simulations of plant water status and drought-induced mortality at the ecosystem level. Geosci. Model Dev. 15, 5593–5626.

79. Sanchez-Martinez, P., Mencuccini, M., Garćıa-Valdés, R., Hammond, W.M., Serra-Diaz, J.M., Guo, W.Y., Segovia, R.A., Dexter, K.G., Svenning, J.C., Allen, C., Martínez-Vilalta, J., 2023. Increased hydraulic risk in assemblages of woody plant species predicts spatial patterns of drought-induced mortality. Nat. Ecol. Evol. 7, 1620–1632. 10.1038/s41559-023-02180-z.

80. Shaw, R.G., Etterson, J.R., 2012. Rapid climate change and the rate of adaptation: insight from experimental quantitative genetics. New Phytol. 195, 752–765. 10.1111/j.1469-8137.2012.04230.x.

81. Sjöman, H., Hirons, A.D., Bassuk, N.L., 2018. Improving confidence in tree species selection for challenging urban sites: a role for leaf turgor loss. Urban Ecosyst. 21, 1171–1188. 10.1007/s11252-018-0791-5.

82. Skelton, R.P., Anderegg, L.D.L., Diaz, J., Kling, M.M., Papper, P., Lamarque, L.J., Delzon, S., Dawson, T.E., Ackerly, D.D., 2021. Evolutionary relationships between drought-related traits and climate shape large hydraulic safety margins in western North American oaks. Proc. Natl. Acad. Sci. U.S.A. 118. 10.1073/pnas.2008987118.

83. Soubeyroux, J.M., Bernus, S., Corre, L., Drouin, A., Dubuisson, B., Etchevers, P., Gouget, V., Josse, P., Kerdoncuff, M., Samacoits, R., Tocquer, F., 2020. Les nouvelles projections climatiques de ŕeférence DRIAS 2020 pour la métropole. Météo France, 97. https://www.drias-climat.fr/.

84. Sánchez-Gómez, D., Aranda, I., 2024. Unveiling intra-population functional variability patterns in a European beech (Fagus sylvatica L.) population from the southern range edge: drought resistance, post-drought recovery, and phenotypic plasticity. Tree Physiol. 44. 10.1093/treephys/tpae107.

85. Tichý, L., Axmanová, I., Dengler, J., Guarino, R., Jansen, F., Midolo, G., Nobis, M.P., Van Meerbeek, K., Ácíc, S., Attorre, F., Bergmeier, E., Biurrun, I., Bonari, G., Bruelheide, H., Campos, J.A., C^̌^arni, A., Chiarucci, A., Cuk, M., Csterevska, R., Didukh, Y., Díťe, D., Díťe, Z., Dziuba, T., Fanelli, G., Fernández-Pascual, E., Garbolino, E., Gavilán, R.G., Gégout, J., Graf, U., Güler, B., Hájek, M., Hennekens, S.M., Jandt, U., Jsková, A., Jiménez-Alfaro, B., Julve, P., Kambach, S., Karger, D.N., Karrer, G., Kavgacı, A., Knollová, I., Kuzemko, A., Küzmc, F., Landucci, F., Lengyel, A., Lenoir, J., Marcenò, C., Moeslund, J.E., Novák, P., PérezHaase, A., Peterka, T., Pielech, R., Pignatti, A., Rsomavcius, V., Rūsiņa, S., Saatkamp, A., S^̌^ilc, U., S^̌^kvorc, v., Theurillat, J., Wohlgemuth, T., Chytrý, M., 2023. Ellenberg-type indicator values for European vascular plant species. J. Veg. Sci. 34. 10.1111/jvs.13168.

86. Tyree, M.T., Hammel, H.T., 1972. The measurement of the turgor pressure and the water relations of plants by the pressure-bomb technique. J. Exp. Bot. 23, 267–282. 10.1093/jxb/23.1.267.

87. Urli, M., Porté, A.J., Cochard, H., Guengant, Y., Burlett, R., Delzon, S., 2013. Xylem embolism threshold for catastrophic hydraulic failure in angiosperm trees. Tree Physiol. 33, 672–683. 10.1093/treephys/tpt030.

88. Valladares, F., Śanchez-Gómez, D., 2006. Ecophysiological traits associated with drought in Mediterranean tree seedlings: Individual responses versus interspecific trends in eleven species. Plant Biol. 8, 688–697. 10.1055/s-2006-924107.

89. Van Genuchten, M., 1980. A closed-form equation for predicting the hydraulic conductivity of unsaturated soils1. Soil. Sci. Soc. Am. J. 44. 10.2136/sssaj1980.03615995004400050002x.

90. Zhu, S.D., Chen, Y.J., Ye, Q., He, P.C., Liu, H., Li, R.H., Fu, P.L., Jiang, G.F., Cao, K.F., 2018. Leaf turgor loss point is correlated with drought tolerance and leaf carbon economics traits. Tree Physiol. 38, 658–663. 10.1093/treephys/tpy013.

91. Zomer, R., Xu, J., Trabucco, A., 2022. Version 3 of the global aridity index and potential evapotranspiration database. Sci. Data 9. 10.1038/s41597-022-01493-1.

