## Supplementary Data for "Beyond Proxies: Towards ecophysiological indicators of drought resistance for forest management"

### Sampled trees

**Table 1.** *Abies* species sampled in the two experimental setups. First setup was planted in 1970 with five year seedlings (Fady et al. 2024). Second setup was planted in 1977 with four year seedlings (Fady et al. 2023). All trees were spaced two meters apart. Setups are in the same forest, one kilometer apart. Survival refers to average species survival. Area of origin of the seeds and its altitude is indicated, along with altitudes found for all species' range area.

| Species | Setup | Provenances | Site altitude (m) (species' range) | Survival |
|---|---|---|---|---|
| *A. borisii-regis* | First | Greece - Kerasini (10) | 1410 (600-2200) | 26% |
| *A. bornmuelleriana* | Second | Türkiye - Ortadil Dokurcun (7) and Aladag (3) | 1321 and 944 (0-2000) | 49% |
| *A. cephalonica* | First | Greece - Xerovounia (10) | 1455 (600-2200) | 26% |
| *A. cilicica* | Second | Türkiye - Hartlap (9) / Syria - Alaouites (1) | 677 and 953 (1000 - 2100) | 32% |
| *A. concolor* | Second | USA - Colorado 1 (3) and Colorado 2 (7) | 59 and 357 (900-3400) | 32% |
| *A. nordmanniana* | Second | Türkiye - Karanlik Mese (8) / Georgia - Tbilissi (2) | 843 and 697 (400-2100) | 49% |
| *A. numidica* | First & Second | Algeria - Babor (10) | 1750 (1850-2000) | NA & 50% |
| *A. pinsapo marocana* | Second | Maroc (10) | 1561 (1400-2000) | 41% |

**Table 2.** Source of trait values for *Abies alba*. URFM : Research Unit Ecology of Mediterranean Forests of the National Research Institute for Agriculture, Food and Environment.

| Trait | Value | Date | Source | Origin of sampled trees |
|---|---|---|---|---|
| TLP | -2.70 $MPa$ | July 2019 | Kunert and Tomaskova (2020) | Fürth, Middle Frankonia, Germany |
| $P_{50}$ | -3.50 $MPa$ | February 2020 | H. Cochard - projet Sap-In | La Courtine, Creuse, France |
| S | 133.2 $\%MPa^{-1}$ | February 2020 | H. Cochard - Sap-In | La Courtine, Creuse, France |
| $g_{min}$ | 1.66 $mmol.m^{-2}.s^{-1}$ | November 2022 | S. Herbette | Neuville, Puy-de-Dôme, France |
| $\epsilon$ | 5.89 $MPa$ | September 2024 | URFM | Ventoux, Vaucluse, France |
| $\pi_0$ | -1.22 $MPa$ | September 2024 | URFM | Ventoux, Vaucluse, France |
| $a_f$ | 0.69 | September 2024 | URFM | Ventoux, Vaucluse, France |
| LDMC | 505.8 $g.g^{-1}$ | September 2024 | URFM | Ventoux, Vaucluse, France |
| LMA | 181.9 $g.m_{leaf}^{-2}$ | September 2024 | URFM | Ventoux, Vaucluse, France |

### Description of "Le Treps" experimental setups

The first experimental setup is composed of eight *Abies* species presented in table 3 They were planted in December 1970. The setup aims at comparing *Abies cephalonica* provenances. Individuals that did not survive were refilled with *Cedrus atlantica*. Setup is described in Fady et al. (2024).

**Table 3.** First setup composition

| Species | Provenances | Percent survival |
|---|---|---|
| *A. alba* | 2 | 0% |
| *A. bornmuelleriana* | 1 | NA |
| *A. cephalonica* | 12 | 26% |
| *A. concolor* | 1 | 17% |
| *A. magnifica* | 1 | NA |
| *A. nordmanniana* | 3 | 30% |
| *A. numidica* | 1 | NA |
| *A. pinsapo pinsapo* | 1 | 16% |
| *C. atlantica* | 1 | 43% |

The second experimental setup is composed of eight *Abies* species. They were planted in February 1977. The setup aims at comparing different *Abies* species. They are presented in table 4. Individuals that did not survive were refilled with *Cedrus atlantica*. Setup is described in Fady et al. (2023).

**Table 4.** Second setup composition

| Species | Provenances | Percent survival |
|---|---|---|
| *A. bornmuelleriana* | 3 | 49% |
| *A. cephalonica* | 1 | 67% |
| *A. cilicica* | 2 | 32% |
| *A. concolor* | 2 | 32% |
| *A. equi-trojani* | 1 | 0% |
| *A. nordmanniana* | 3 | 49% |
| *A. numidica* | 1 | 50% |
| *A. pinsapo marocana* | 1 | 41% |
| *C. atlantica* | 8 | NA |

### Operational indicators

**Table 5.** Data compilation for *Abies* species. Species in **bold** refer to "**Mean**" values extracted for the entire range area of the species. Species in *italic* refer to original location of *"sampled"* seeds, used for the common garden experiment of Le Treps. Bioclimatic variables (Fick and Hijmans 2017), aridity and potential evapotranspiration indexes (Zomer et al. 2022) were extracted according to European Forest Genetic Resources Programme's range area (euforgen.org, last accessed in August 2023) and according to Global Biodiversity Information Facility (gbif.org, last accessed in August 2023) for *Abies concolor* which is not a European species. 5% quantile, mean and 95% quantile are displayed for average species values.

| Species | Bioclimatic indicators - Q5 - **mean** - Q95 | | | | | | |
|---|---|---|---|---|---|---|---|
| **Mean** *Sampled* | Annual mean temperature (°C) - bio1 | Mean temperature of warmest quarter (°C) - bio10 | Annual precipitation (mm) - bio12 | Precipitation seasonality (mm) - bio15 | Precipitation of warmest quarter (mm) - bio18 | Aridity Index - AI | Potential evapotranspiration ET0 |
| ***A. alba*** | 3.7 - **7.5** - 10.6 | 11.9 - **16.1** - 19.0 | 544 - **897** - 1423 | 13.2 - **29.6** - 47.2 | 160 - **275** - 459 | 6096 - **10120** - 16948 | 752 - **900** - 1164 |
| ***A. borisii-regis*** | 7.0 - **11.3** - 15.3 | 15.1 - **20.0** - 24.9 | 440 - **753** - 1288 | 21.2 - **35.4** - 55.0 | 59 - **103** - 140 | 3076 - **5960** - 10562 | 1099 - **1244** - 1408 |
| *A. borisii-regis* | 9.0 | 17.1 | 860 | 40.9 | 101 | 7186 | 1196 |
| ***A. bornmuelleriana*** | 5.8 - **8.5** - 11.8 | 14.3 - **17.0** - 20.1 | 550 - **653** - 777 | 19.5 - **29.4** - 38.8 | 105 - **128** - 156 | 4349 - **5471** - 6626 | 1102 - **1198** - 1305 |
| *A. bornmuelleriana* | 7.8 | 15.7 | 617 | 30.6 | 112 | 5268 | 1171 |
| *A. bornmuelleriana* | 9.4 | 17.5 | 551 | 33.3 | 98 | 4461 | 1234 |
| ***A. cephalonica*** | 6.8 - **11.1** - 15.1 | 14.9 - **19.4** - 23.4 | 541 - **817** - 987 | 29.9 - **51.9** - 68.0 | 38 - **75** - 123 | 3796 - **6363** - 8611 | 1103 - **1292** - 1463 |
| *A. cephalonica* | 9.6 | 17.6 | 923 | 62.0 | 64 | 7093 | 1301 |
| ***A. cilicica*** | 5.6 - **10.2** - 15.0 | 15.0 - **19.5** - 23.8 | 507 - **638** - 810 | 50.5 - **61.8** - 79.8 | 23 - **42** - 57 | 3413 - **4205** - 5123 | 1351 - **1514** - 1681 |
| *A. cilicica* | 15.6 | 25.6 | 707 | 74.6 | 18 | 4019 | 1758 |
| *A. cilicica* | 15.7 | 23.6 | 1229 | 88.5 | 27 | 7208 | 1704 |
| ***A. concolor*** | 4.9 - **8.4** - 12.7 | 14.6 - **17.6** - 22.4 | 337 - **496** - 772 | 18.4 - **48.8** - 77.7 | 64 - **146** - 287 | 1657 - **2794** - 4181 | 1483 - **1819** - 2227 |
| *A. concolor* | 16.3 | 24.3 | 211 | 80.7 | 3 | 1092 | 1931 |
| *A. concolor* | 17.9 | 26.7 | 384 | 82.9 | 5 | 1634 | 2349 |
| ***A. nordmanniana*** | 0.9 - **7.3** - 12.1 | 9.8 - **16.0** - 20.8 | 531 - **955** - 1583 | 14.6 - **26.4** - 41.8 | 76 - **220** - 375 | 4185 - **9072** - 16028 | 873 - **1105** - 1310 |
| *A. nordmanniana* | 12.0 | 22.0 | 554 | 43.9 | 180 | 4560 | 1214 |
| *A. nordmanniana* | 11.4 | 20.4 | 868 | 22.7 | 184 | 6951 | 1248 |
| ***A. numidica*** | 9.7 - **11.3** - 13.2 | 19.1 - **20.7** - 22.5 | 827 - **921** - 1023 | 54.5 - **55.5** - 57.4 | 53 - **60** - 68 | 5126 - **5881** - 6724 | 1524 - **1568** - 1615 |
| *A. numidica* | 9.8 | 18.9 | 1020 | 55.0 | 67 | 6974 | 1462 |
| ***A. pinsapo marocana*** | 10.1 - **12.8** - 16.4 | 18.0 - **20.6** - 24.1 | 890 - **1018** - 1113 | 73.3 - **78.5** - 83.3 | 15 - **21** - 29 | 5145 - **6438** - 7139 | 1478 - **1586** - 1731 |
| *A. pinsapo marocana* | 13.9 | 22.8 | 484 | 57.5 | 28 | 2480 | 1951 |
| **Treps site** | **12.3** | **19.4** | **844** | **38.7** | **128** | **6447** | **1309** |

**Table 6.** Drought-resistance indicators for *Abies* species : ClimEssences mature tree's resistance to extreme drought (RMT AFORCE 2021), resistance to desiccation of excised branches (Aussenac 1980), xericity index (Rameau et al. 2016) (Values range from H to XXX : hygrophile (H) to hyperxerophile (XXX); *m : mesophile; x : xerocline ; X : mesoxerophile ; XX : xerophile* ), Ellenberg-type indicator (Tichý et al. 2023). For ClimEssences and Aussenac, D corresponds to poorly resistant to drought and A very resistant. For Ellenberg, the higher the number, the more resistant is the species.

| | Drought resistance indicators | | | |
|---|---|---|---|---|
| Species | ClimEssences' index (RMT AFORCE 2021) A-D | Resistance to desiccation (Aussenac 1980) A-D | Xericity index (Rameau et al. 2016) H - XXX | Ellenberg (Tichý et al. 2023) 1 - 12 |
| *A. alba* | D | D | m to X- | 6.1 |
| *A. borisii-regis* | na | *na* | XX | 4.9 |
| *A. bornmuelleriana* | B | *na* | XX- | *na* |
| *A. cephalonica* | A | A | XX | 4 |
| *A. cilicica* | B | A | *na* | *na* |
| *A. concolor* | C | A | x to X | *na* |
| *A. nordmanniana* | B | C | XX- | 5 |
| *A. numidica* | A | B | XX- | *na* |
| *A. pinsapo marocana* | A | C | XX | 4 |

### Analysis of Variance

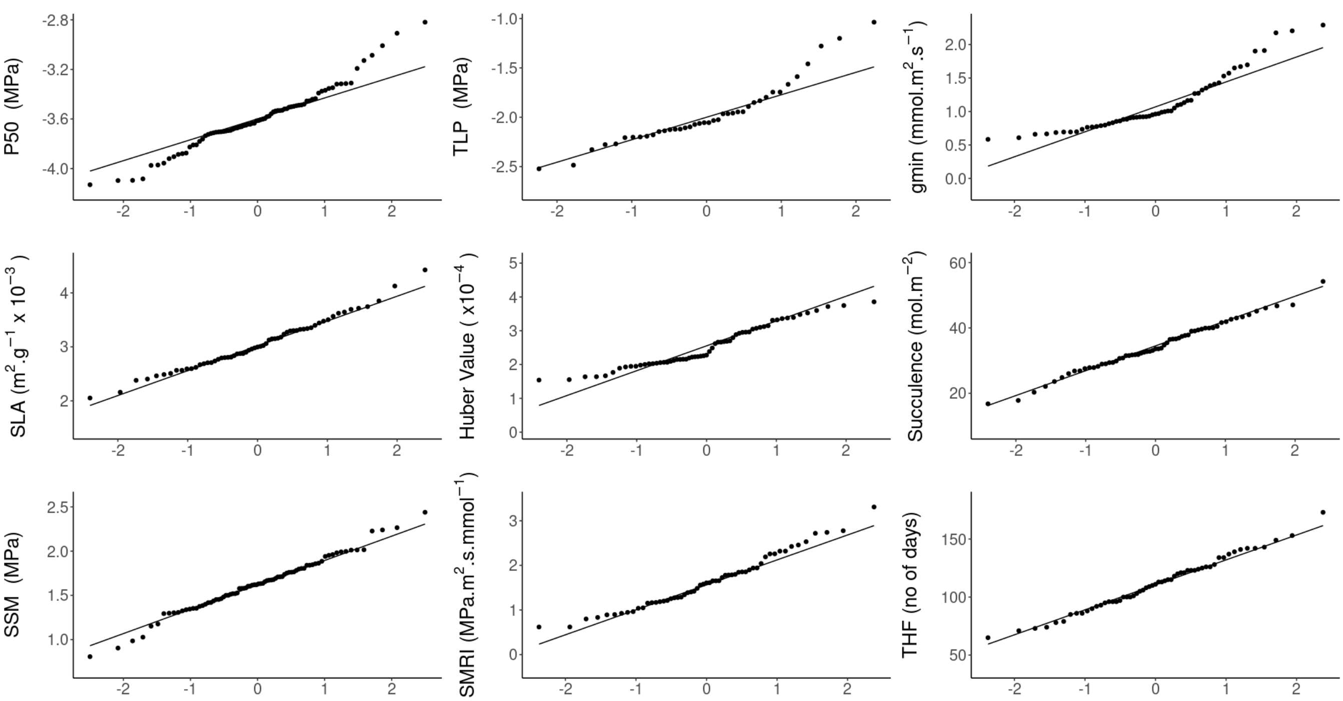


Fig. 1: QQPlot for ANOVA analysis

### Time to Hydraulic Failure regressed on traits

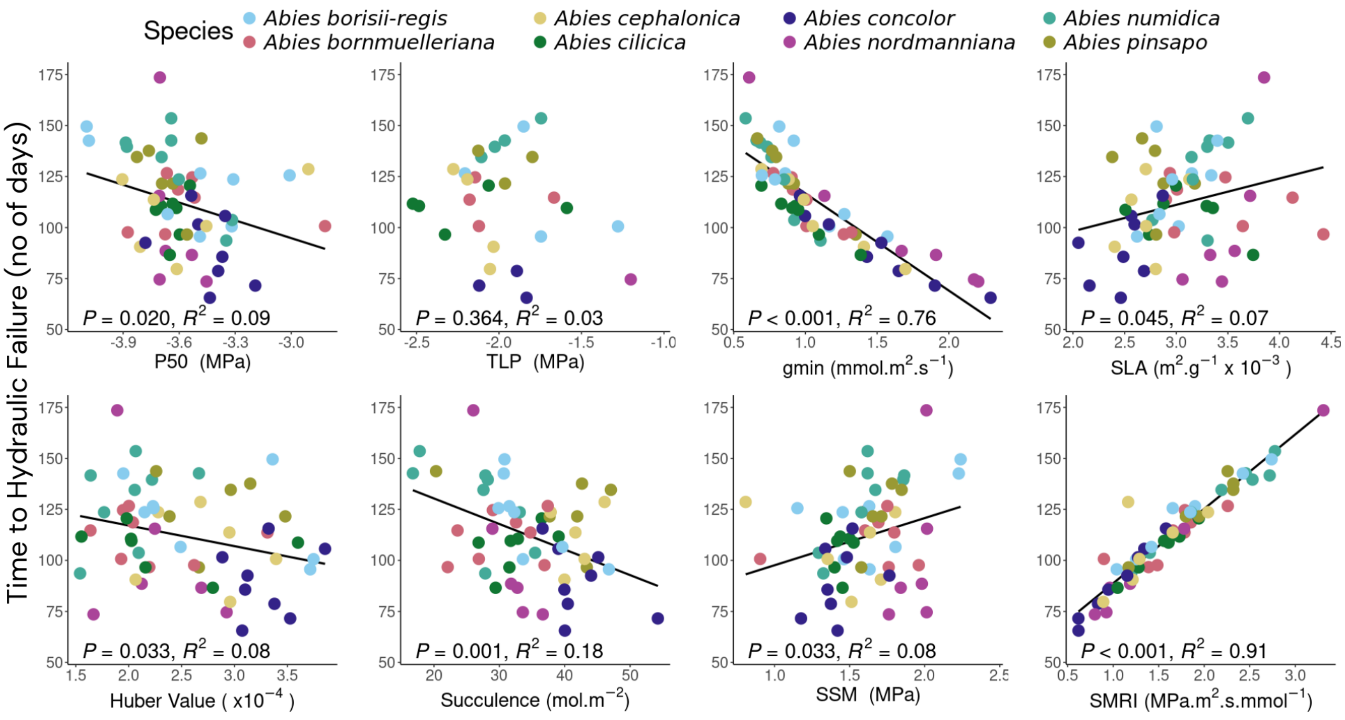


Fig. 2: Regression of Time to Hydraulic Failure on different traits. $P_{50}$: water potential at 50% loss of conductivity, TLP: water potential at turgor loss point, $g_{min}$: minimal water losses after stomatal closure, SLA: Specific Leaf Area, Huber Value, Succulence, SSM: Stomatal Safety Margin, SMRI: Stomatal Margin Retention Index.

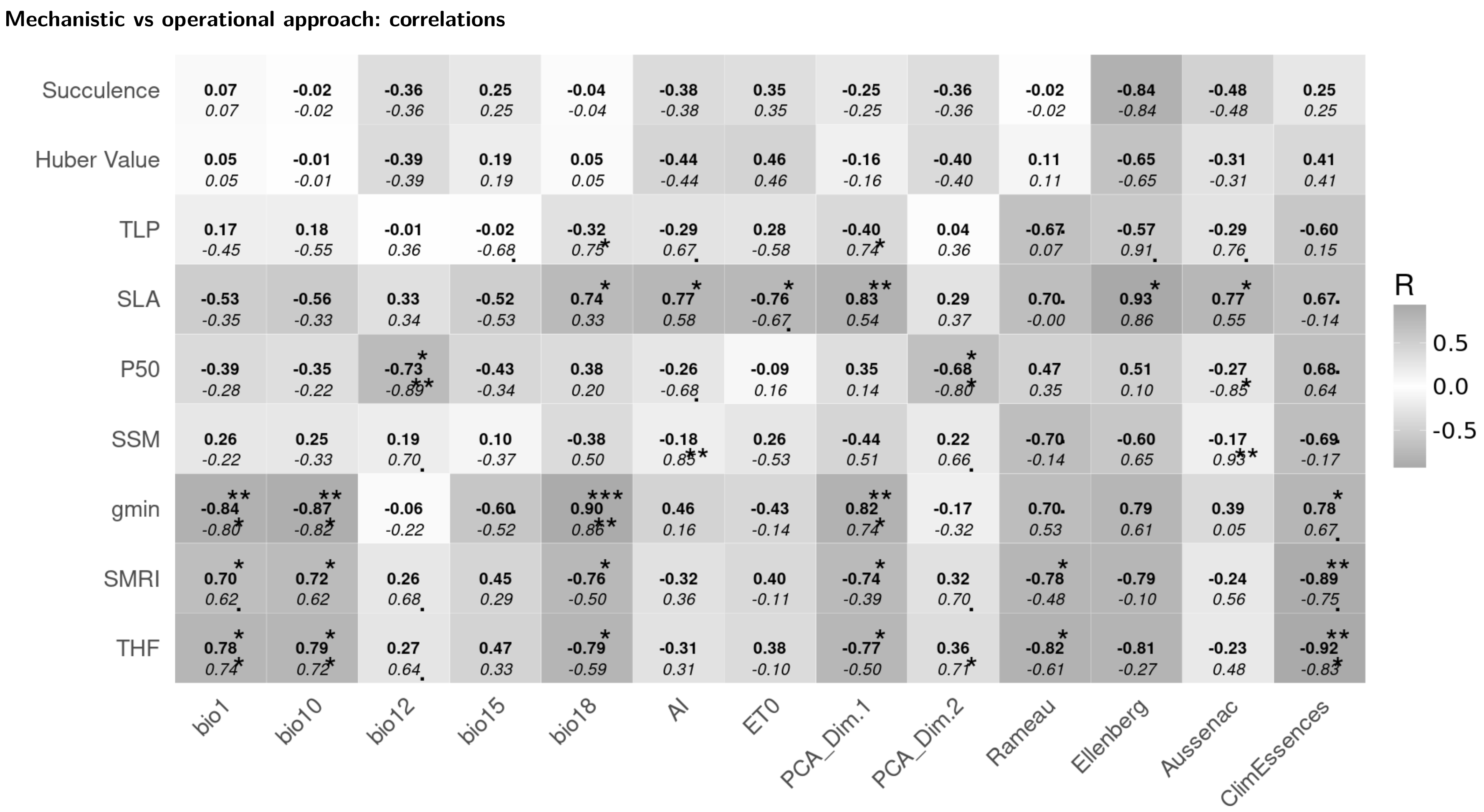


**Fig. 3.** Pearson correlation coefficients of mechanistic traits versus "operational" indicators using mean species values. The above value in **bold** is the correlation when *Abies alba* is included in the dataset. The lower value in *italic* is the correlation when *Abies alba* is removed from the dataset. $p_{values}$ are indicated by the following symbology: $*** < 0.001$ ; $** < 0.05$ ; $* < 0.01$, $. < 0.1$. Succulence, Huber Value, TLP: water potential at turgor loss point, SLA: Specific Leaf Area, $P_{50}$: water potential at 50% loss of conductivity, SSM: Stomatal Safety Margin, $g_{min}$: minimal water losses after stomatal closure, SMRI: Stomatal Margin Retention Index, THF: Time to Hydraulic Failure.
